# Patient-derived colon epithelial organoids reveal lipid-related metabolic dysfunction in pediatric ulcerative colitis

**DOI:** 10.1101/2024.08.22.609271

**Authors:** Babajide A. Ojo, Lyong Heo, Sejal R. Fox, Amanda Waddell, Maria E. Moreno-Fernandez, Marielle Gibson, Tracy Tran, Ashley L. Dunn, Essam I.A Elknawy, Neetu Saini, Javier A. López-Rivera, Senad Divanovic, Vinicio A. de Jesus Perez, Michael J. Rosen

**Affiliations:** Division of Pediatric Gastroenterology, Hepatology, and Nutrition, Stanford University School of Medicine; Stanford, CA, USA; Center for IBD and Celiac Disease, Stanford Medicine Children’s Health; Palo Alto, CA, USA; Stanford Center for Genomics and Personalized Medicine, Stanford University; Palo Alto, CA, USA; Applied Gene and Cell Therapy Center, Cincinnati Children’s Hospital Medical Center; Cincinnati, OH, USA; Department of Pediatrics, University of Cincinnati College of Medicine; Cincinnati, OH, USA; Division of Immunobiology, Cincinnati Children’s Hospital Medical Center; Cincinnati, OH, USA; Division of Gastroenterology, Hepatology, and Nutrition, Cincinnati Children’s Hospital Medical Center; Cincinnati, OH, USA; Division of Pediatric Hematology, Oncology, Stem Cell Transplantation and Regenerative Medicine, Stanford University School of Medicine; Palo Alto, CA, USA; Center for Inflammation and Tolerance, Cincinnati Children’s Hospital Medical Center; Cincinnati, OH, USA; Division of Pulmonary and Critical Care Medicine, Department of Medicine, Stanford University; Stanford, CA, USA

**Keywords:** Ulcerative Colitis, Colon Epithelium, Patient-Derived Organoids, Epithelial Metabolism, Inflammatory Bowel Diseases, Oxidative Phosphorylation

## Abstract

**Background & Aims:** Ulcerative colitis (UC) is associated with epithelial metabolic derangements which exacerbate gut inflammation. Patient-derived organoids recapitulate complexities of the parent tissue in health and disease; however, whether colon organoids (colonoids) model metabolic impairments in the pediatric UC epithelium is unclear. This study determined the functional metabolic differences in the colon epithelia using epithelial colonoids from pediatric patients.

**Methods:** We developed biopsy-derived colonoids from pediatric patients with endoscopically active UC, inactive UC, and those without endoscopic or histologic evidence of colon inflammation (non-IBD controls). We extensively interrogated metabolic dysregulation through extracellular flux analyses and tested potential therapies that recapitulate or ameliorate such metabolic dysfunction.

**Results:** Epithelial colonoids from active UC patients exhibit elevated oxygen consumption and proton leak supported by enhanced glycolytic capacity and dysregulated lipid metabolism. The hypermetabolic features in active UC colonoids were associated with increased cellular stress and chemokine secretion, specifically during differentiation. Transcriptomic and pathway analyses indicated a role for PPAR-α in lipid-induced hypermetabolism in active UC colonoids, which was validated by PPAR-α activation in non-IBD colonoids. Accordingly, limiting neutral lipid accumulation in active UC colonoids through pharmacological inhibition of PPAR-α induced a metabolic shift towards glucose consumption, suppressed hypermetabolism and chemokine secretion, and improved cellular stress markers. Control and inactive UC colonoids had similar metabolic and transcriptomic profiles.

**Conclusions:** Our pediatric colonoids revealed significant lipid-related metabolic dysregulation in the pediatric UC epithelium that may be alleviated by PPAR-α inhibition. This study supports the advancement of colonoids as a preclinical human model for testing epithelial-directed therapies against such metabolic dysfunction.

**What You Need to Know:** *Background and Context:* Colon mucosa healing in pediatric UC requires reinstating normal epithelial function but a lack of human preclinical models of the diseased epithelium hinders the development of epithelial-directed interventions.

**New Findings:** Using colon biopsy-derived epithelial organoids, samples from pediatric patients with active UC show hyperactive metabolic function largely driven by enhanced lipid metabolism. Pharmacologic inhibition of lipid metabolism alleviates metabolic dysfunction, cellular stress, and chemokine production.

**Limitations:** Though our epithelial colon organoids from active UC patients show targetable metabolic and molecular features from non-IBD controls, they were cultured under sterile conditions, which may not fully capture any potential real-time contributions of the complex inflammatory milieu typically present in the disease.

**Clinical Research Relevance:** Current therapies for pediatric UC mainly target the immune system despite the need for epithelial healing to sustain remission. We identified a pharmacologic target that regulates epithelial metabolism and can be developed for epithelial-directed therapy in UC.

*Basic Research Relevance:* Pediatric UC patient tissue adult stem cell-derived colon epithelial organoids retain disease-associated metabolic pathology and can serve as preclinical human models of disease. Excess reliance on lipids as an energy source leads to oxidative and inflammatory dysfunction in pediatric UC colon organoids. **Preprint:** This manuscript is currently on bioRxiv. doi: https://doi.org/10.1101/2024.08.22.609271 **Lay Summary:** Using patient tissue-derived colon epithelial organoids, the investigators identified epithelial metabolic dysfunction and inflammation in pediatric ulcerative colitis that can be alleviated by PPAR-a inhibition.

## Main Text

The restoration of normal intestinal epithelial function is critical for colon mucosa healing in inflammatory bowel diseases (IBD); ^1,2^ however, current therapies predominantly target the immune system. As such, only 20-30% of patients achieve mucosal healing with today’s therapies.^3–6^ Ulcerative colitis (UC) is an IBD subtype characterized by the continuous chronic inflammation of the colon mucosa, extending proximally from the rectum.^7^ Pediatric UC is marked by more severe and extensive disease than adult UC.^7^ Achieving mucosa healing is one of the best predictors of sustained disease remission.^8^ Studies that enhance our understanding of the human intestinal epithelial function in health and disease may advance epithelial-directed therapies in pediatric UC.

Systemic and local dysregulation of metabolism is a hallmark of inflammation in human IBD.^9,10^ Direct investigation of the inflamed human colon tissue in UC has proposed several metabolic impairments in UC, including impaired mitochondrial respiratory chain complex activity, enhanced epithelial mitochondria fission, deficient butyrate metabolism, and reduced epithelial mitochondrial membrane potential.^11–14^ This may be critical to the course of disease since mitochondrial activity regulates several aspects of epithelial cellular homeostasis, including the maintenance of energetic balance, adequate reactive oxygen species (ROS) production, and regulating pro-inflammatory responses.^15^ However, the cellular mechanisms underlying UC epithelial metabolic dysfunction are not known. Such a deeper understanding may reveal novel epithelial targets to promote mucosal healing.

The development of epithelial-directed therapies in UC is hindered by a lack of human preclinical models of the diseased epithelium. Over the last decade, advances in organoid biology have grown their promise as human pre-clinical models of disease.^16^ In particular, patient-derived adult stem cell organoids are capable of self-organizing into structural and functional models that may reflect the cell and molecular complexities of parent tissues.^17^ In UC, patient-derived colon organoids (colonoids) may reflect disease-specific disparities compared to healthy colonoids including alterations in transcription, epigenetics, and somatic gene mutations.^18–22^ Despite the promise of intestinal organoids in understanding human intestinal diseases, they have not been used to interrogate the metabolic nature of disease in pediatric UC. In this study, we used mucosal biopsy-derived epithelial colonoids from non-IBD, active, and inactive pediatric UC patients to understand the functional metabolic features of the UC epithelium. We further assessed the targetable molecular features under which metabolic dysregulation in active UC can be ameliorated.

## MATERIALS AND METHODS

### Human Study Approval and Informed Consent

Pediatric patient recruitment occurred at Cincinnati Children’s Hospital Medical Center, Cincinnati, Ohio, and Lucile Packard Children’s Hospital Stanford, Palo Alto, California. Patient data and biopsy collection followed approved protocols by the institutional review boards at each institution. Four to eight colon biopsies (2–5 mm^3^) from patients with active UC, inactive UC, and non-IBD controls were collected into ice-cold DMEM/F12 (Corning 10-092-CV) + 50 µg/mL Normocin (InvivoGen #ant-nr) during clinically-indicated lower endoscopy and transported on ice to the lab for immediate processing for spheroid generation. Patient demographics and clinical data are presented in Supplementary Table 1.

### Biopsy processing and organoid culture

Organoids were generated from fresh biopsies as detailed in Supplementary Methods.

### Spheroids processing and plating for analysis

After 5-7 days after passaging with TrypLE Express, Matrigel domes with matured spheroids were dislodged with ice-cold PBS as previously described and filtered with a 100 µm strainer (pluriSelect). Filtrates were pelleted and washed once in OWM, manually counted, and replated in domes containing a 1:1 mix of Matrigel and OWM (8-15 spheroids per µL depending on the analysis) and maintained in Intesticult human organoid differentiation medium, OGM, (STEMCELL Technologies, #06010). For all analyses requiring differentiation into colonoids, replated spheroids were allowed to acclimatize overnight in OGM before switching to Intesticult human organoid differentiation medium, ODM, (STEMCELL Technologies, #100-0214) for the indicated differentiation durations with ODM changed every 2 days until analysis. See Supplementary Methods for details.

### Extracellular Flux Analyses

The XF Seahorse MitoStress Test (Agilent) was used to measure oxygen consumption rates (OCR), at baseline, and after the injection of mitochondrial inhibitors. Each spheroid line was replated in quadruplicates as described previously in 15 µL matrigel domes (8 spheroids/µL) containing a 1:1 mix of Matrigel and OWM into Seahorse XF96 cell culture plates (Agilent #101085-004) and cultured overnight in 150 µL OGM. The four edges of the plate served as blank with only a 1:1 mix of Matrigel and OWM. For some experiments, 50 µL of OGM was removed after 24 hrs into an empty 96-well plate and stored at -80°C for later measurements of LDH activity. Where differentiation was needed, spheroids were washed once with PBS on the following day and differentiated in 200 µL ODM into colonoids for the indicated number of days in the figure legends.

On the day of the assay, ODM was removed from colonoids, washed once with 200 µL PBS, and replaced with 150 µL DMEM assay medium (Agilent, #103334-100), pH 7.4 containing 10 mM glucose, 2 mM glutamine, and 1 mM pyruvate (all from Agilent). Plates were incubated in a non-CO_2_ incubator for 1 hr and OCR was measured at baseline and after a sequential injection of oligomycin (final concentration, 4µM), FCCP (final concentration, 2 µM), and rotenone/antimycin (final concentration, 100 nM/1 µM) using the XF96^e^ Extracellular Flux assay kits on the XF96^e^ analyzer. Data normalization was carried out with DNA content as previously described for organoids^23^ and detailed in Supplementary Methods.

See Supplementary Methods for details on all other measurements.

### Statistical analyses

Eight patient-derived colonoids were generated per diagnosis (control, active UC, inactive UC). For all data, symbols in figures represent the average measures of individual colonoid lines used for each assay. Analysis of two groups was conducted with a two-sided Mann-Whitney test or unpaired T-test. Wherever colonoid lines from the same diagnosis were treated with a Fccp, Fenofibrate, or GW6471, statistical analyses were done with a two-sided paired T-test or Wilcoxon signed-rank test. Two-way ANOVA with Tukey’s *post hoc* test was used for etomoxir experiments and those collected over 5 days (day 2 and day 5). Statistical analysis was conducted in Prism (v.10.0), and *P* values < 0.05 were considered statistically significant. For differentially expressed genes from the RNA-seq data, transcripts with Log_2_FC ≥ ±1, and FDR ≤ 0.1 were considered statistically significant.

## RESULTS

### Derivation of pediatric colon organoids

We developed organoids from rectal biopsies obtained from pediatric patients (n=24) undergoing clinically indicated colonoscopies (Supplementary Figure 1*A*, see Supplementary Table 1 for patient data). We observed enhanced recovery of matured non-differentiated spheroids from biopsies obtained from non-IBD controls by day 7 (Supplementary Figure 1*B*), whereas aUC spheroids required 10-14 days. After the extended growth period, fully established aUC spheroids were morphologically comparable to controls and maintained similarity after multiple passages and a freeze-thaw cycle (Supplementary Figure 1C-D). The timeline and morphology of spheroids generated from inactive UC (iUC) biopsies (patients with endoscopic healing from active disease) were similar to that of biopsies from non-IBD patients.

### Colonoids from active UC patients exhibit impaired differentiation and hypermetabolism

Under non-differentiated conditions, we found no differences in the spheroid diameter between control and aUC spheroids (Supplementary Figure 2*A* and *B*). Accordingly, we observed no metabolic differences between control and aUC spheroids (Supplementary Figure 2*C-E*). Similarly, we analyzed extracellular LDH release into the medium – a marker cellular cytotoxicity – and found no differences in LDH release between control and aUC spheroids (Supplementary Figure 2*F*). These results suggest that control and aUC spheroids are morphologically and metabolically similar in their undifferentiated states.

In contrast, we observed morphologic differences between aUC and control colon organoids under differentiating conditions (hereafter referred to as colonoids) as early as day 3 as more aUC colonoids remained in their cystic morphology compared to the largely budded structures of control colonoids (Figure 1*A* and *B*). Further differentiation up to 7 days resulted in visibly reduced survival in aUC colonoids compared to controls (Supplementary Figure 2*G*). To complement these morphological features, we determined the gene expression of the secretory lineage progenitor, *ATOH1*, the enteroendocrine cell marker, *CHGA*, and the crypt-based goblet cell marker of the colon, *WFDC2*,^24^ in C and aUC before and after 3-day differentiation (i.e spheroids vs colonoids). None of the examined markers were significantly different in the control and aUC spheroids (Figure 1*C*, and Supplementary Figure 2*H-I*). As expected, *ATOH1* and *CHGA* were upregulated in both control and aUC colonoids compared to their respective undifferentiated spheroids (Supplementary Figure 2*H* and *I*). In aUC colonoids, there was a failure to upregulate WFDC2, consistent with the phenomenon of goblet cell depletion in UC (Figure 1*C*).^24^

**Figure 1.**
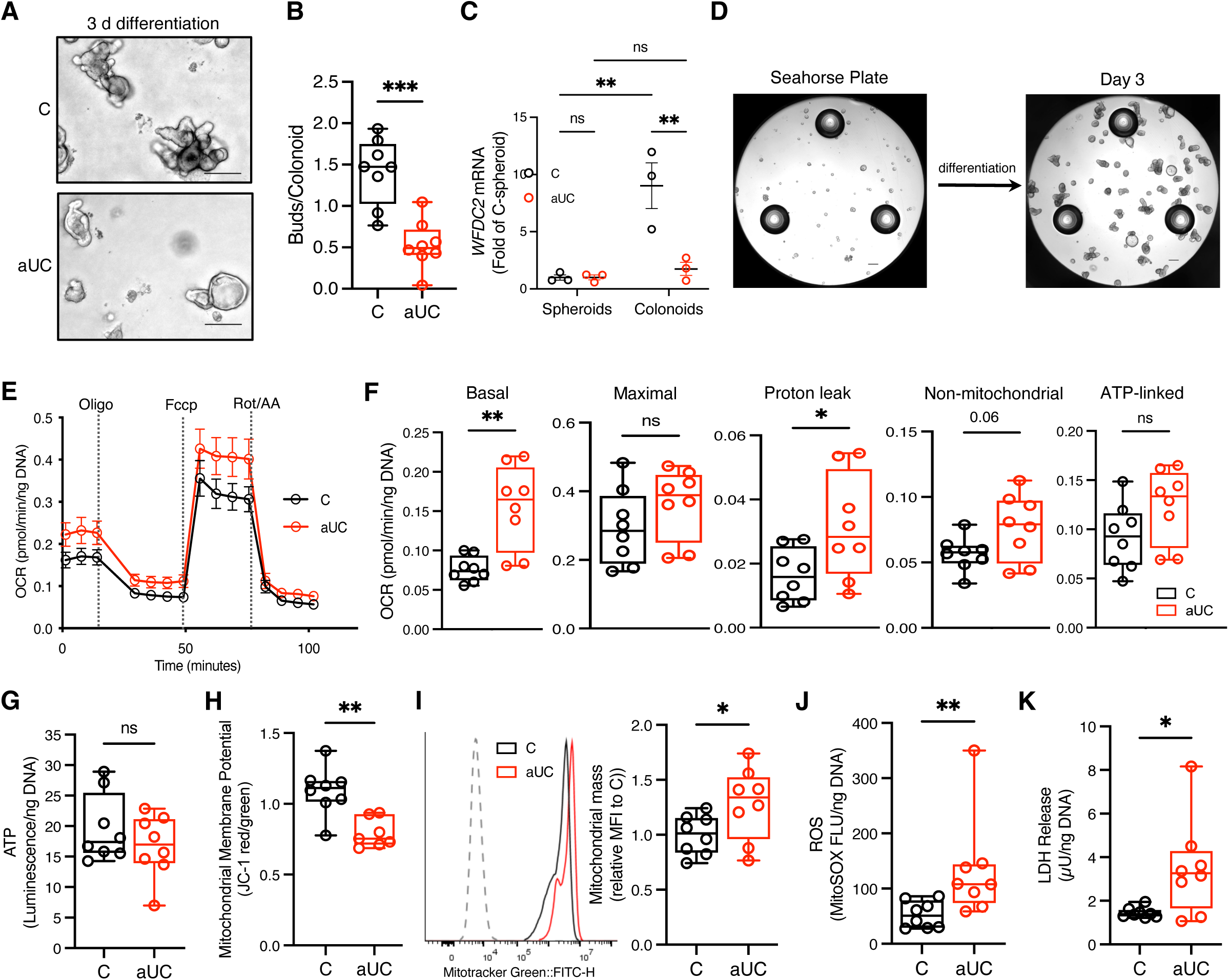
Hypermetabolism and cellular stress in active UC colonoids during differentiation. (**A**) Representative phase contrast images of pediatric control (C) and active (a)UC colonoids after 3-day differentiation. Scale bar, 200 µm. (**B**) Buds/colonoid in colonoids treated as in **A**. (**C**) *WFDC2* mRNA abundance using qRT-PCR in paired spheroids (undifferentiated) and colonoids differentiated for 3 days. (**D**) Phase contrast image of a Seahorse 96-well plate with a representative spheroid line before and after 3-day differentiation. Scale bar, 200 µm. (**E**) Bioenergetic profile of C and aUC colonoids treated as in **A**, and subjected to the Seahorse MitoStress test. (**F**) MitoStress OCR response of C and aUC colonoids. (**G**) Total ATP luminescence assay. (**H**) Colonoids were treated as in **A**, and Mitochondrial membrane potential (MMP) estimation with the JC-1 dye by flow cytometry. (**I**) Mitochondrial mass estimated with Mitotracker green intensity in C and aUC colonoids treated as in **A**. (**J**) Mitochondrial ROS estimates in colonoids cultured as in **A**, with the MitoSOX fluorescence assay. (**K**) LDH activity in C and aUC colonoids after 2 days. *=*P* < 0.05, **=*P* < 0.01, ***=*P* < 0.001. n=3-8 patient colonoid lines/group. For all data, symbols represent the average measures of individual colonoid lines. LDH, lactate dehydrogenase; OCR, oxygen consumption rates; ROS, reactive oxygen species

Following the evidence of impaired differentiation in aUC colonoids, we examined whether the metabolic profiles of control and aUC colonoids diverged during the early stages of differentiation. After day 3 of differentiation, we observed a 122% elevation of basal OCR in aUC colonoids compared to the controls. In addition, we found a 75% significant elevation in median OCR due to proton leak in aUC colonoids compared to control, with a non-significant increase in non-mitochondrial and ATP-linked OCR compared to control colonoids (Figure 1*D*-*F*). Similarly, we found no differences in total cellular ATP between aUC and control colonoids (Figure 1*G*). Colon epithelial cells from treatment-naïve UC patients exhibit lower mitochondrial membrane potential (MMP),^14^ and we observed a similar reduction in MMP in aUC colonoids compared to controls (Figure 1*H*). In line with the hypermetabolic patterns in aUC colonoids, we observed a significant elevation of mitochondrial mass (mtMass) during differentiation compared to controls (Figure 1*I*). Correspondingly, we found significant increases in mitochondrial ROS levels and LDH release in aUC colonoids compared to control, suggesting that hypermetabolism in aUC colonoids is accompanied by enhanced oxidative stress and cytotoxicity (Figure 1*J* and *K*). These data suggest hypermetabolism and metabolic inefficiency in aUC colonoids at the early stages of differentiation which is associated with cellular stress.

Finally, we asked whether colonoids from inactive UC (iUC) patients were metabolically distinct from control colonoids. First, we observed similar morphological and metabolic differences between control and iUC colonoids during differentiation (Supplementary Figure 3*A-E*); however, non-mitochondrial OCR was higher in iUC compared to control colonoids (Supplementary Figure 3*D*). Similar to our observation in aUC colonoids, we observed enhanced ROS in iUC colonoids compared to control colonoids.

### Metabolic uncoupling contributes to hypermetabolism in aUC colonoids

The proton leak and resulting metabolic inefficiency observed in aUC colonoids during differentiation may be indicative of enhanced uncoupling of mitochondrial respiration.^25^ Thus, we hypothesized that endogenous mitochondrial uncoupling proteins would be upregulated in aUC compared to control. Analyses of bulk RNA-seq datasets from the RISK study (GSE117993)^26^ and PROTECT Pediatric IBD inception cohort studies (GSE109142)^14^ showed that *UCP2* is significantly upregulated in the rectal mucosa of pediatric UC patients compared to controls (Supplementary Figure 4*A-C*). Similarly, we observed enhanced UCP2 expression at the mRNA and protein levels in our patient-derived colonoid samples (Supplementary Figure 4*D* and *E*).

To test whether metabolic uncoupling was sufficient to induce the altered colon epithelial differentiation and metabolism observed aUC, we added low-dose Fccp (protonophore that uncouples mitochondrial OXPHOS), into the medium throughout a 3-day differentiation of control colonoids (Supplementary Figure 4*F*). Fccp raised basal OCR and OCR due to maximal respiration, proton leak, and non-mitochondrial respiration, but also significantly increased ATP-linked respiration, suggesting that the healthy colonoids may overcome chemical uncoupling to maintain ATP levels (Supplementary Figure 4*G* and *H*). Still, we observed elevated mtMass and mitochondrial ROS levels in Fccp-treated colonoids compared to controls (Supplementary Figure 4*I* and *J*). In contrast, we did not see an increase in LDH release to suggest uncoupling alone induced cytotoxicity (Supplementary Figure 4*K*). Fccp did not impact colonoid morphology or *WFDC2* gene expression (Supplementary Figure 4*L-N*). These data suggest that upregulation of UCP2 and mitochondrial uncoupling likely contribute to hypermetabolism and oxidative stress in aUC colonoids, but may be insufficient to impair differentiation or induce cytotoxicity.

### Active UC colonoids exhibit an altered transcriptional profile and upregulated chemokine secretion

To understand transcriptional changes that may underly altered metabolism and function in aUC colonoids, we conducted bulk RNA-seq on spheroids and colonoids differentiated for 3 days (n= 8 patient lines per diagnosis, passage 5-7). Only seven genes were significantly upregulated and four were significantly downregulated in aUC compared to control undifferentiated spheroids (Log_2_FC ≥ ±1, FDR P ≤ 0.1, Supplementary Datasheet 1). After 3 days of differentiation, control and aUC colonoids had divergent transcriptomic profiles (Figure 2*A*), in line with their distinct metabolic profiles. Specifically, 112 genes were significantly upregulated and 34 significantly downregulated (Log_2_FC ≥ ±1, FDR P ≤ 0.1; Supplementary Datasheet 1). The top upregulated genes included well-known MHCII-related genes in active UC,^14,27^ *HLA-DRA, HLA-DMB, HLA-DRB, HLA-DMA*; lipid-related genes (*FABP6, ABCA12, and APOL3*); and *CARD6* a member of the caspase recruitment domain family (Figure 2*B*). Top downregulated genes included those involved in the development of cytoskeletal filaments (*NEFH, KRT12*), goblet cells (*ZG16, WFDC2*), and those involved in cellular development and histone maintenance (*GATA2* and *HIST1H4K*, respectively).

**Figure 2.**
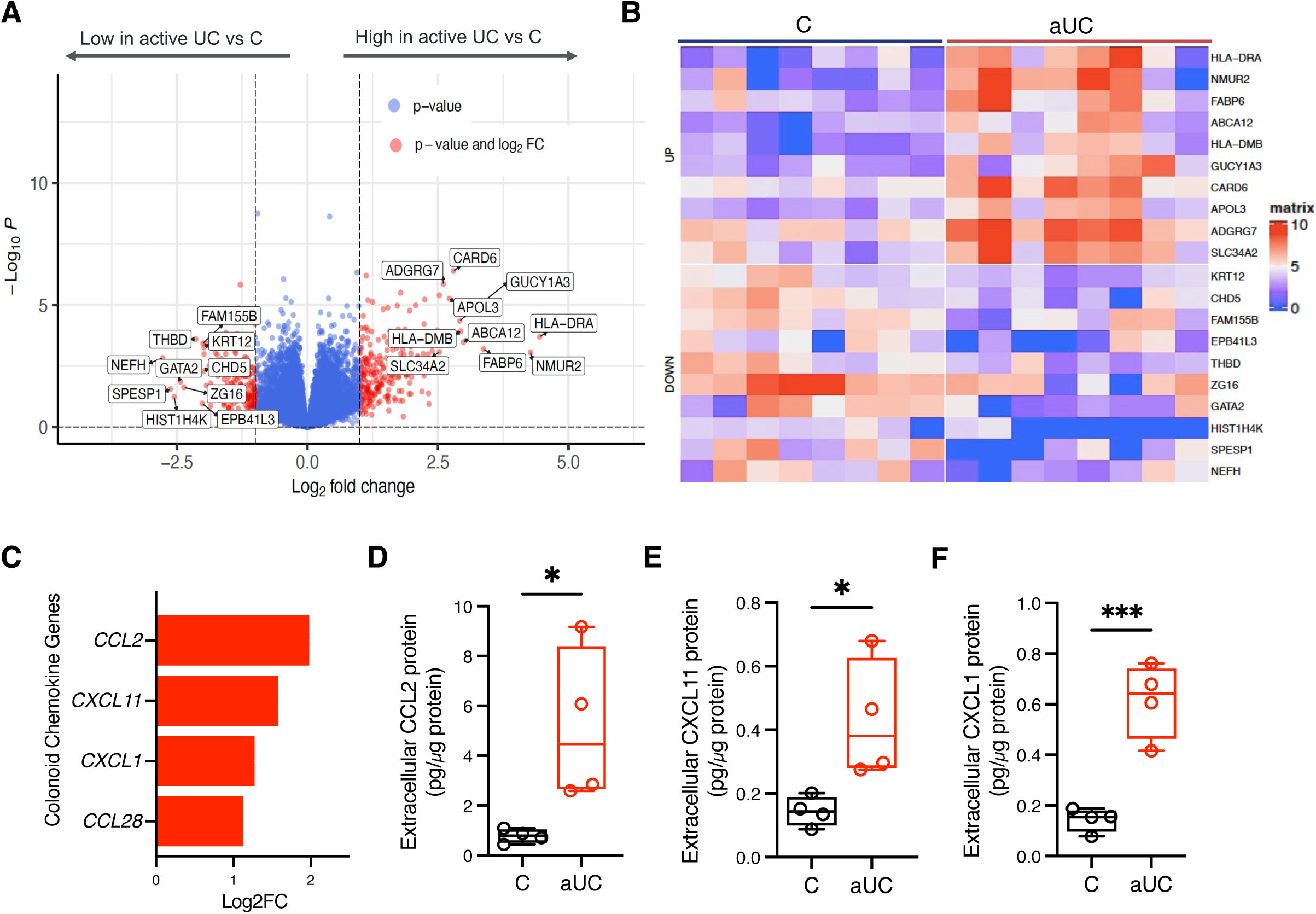
Distinct transcriptome in epithelial colonoids from active UC pediatric patients. Pediatric epithelial colonoids differentiated for 3 days and were subjected to bulk RNA-seq. n=8 patient colonoid line/group. (**A**) Volcano plot of differentially expressed genes between aUC and C samples (Log2FC>1, FDR <0.1). The full list of differentially expressed genes is provided in Supplementary DataSheet 1. (**B**) Heat map of the top 10 dysregulated genes showing the diversity and consistency of expression among each colonoid line. (**C**) Chemokine genes overexpressed in aUC epithelial colonoids compared to C in the bulk RNA-seq data (Log2FC>1, FDR <0.1). (**D**) Extracellular chemokine protein, CCL2, (**E**) CXCL11, and (**F**) CXCL1 after a 3-day differentiation. n=4 patient colonoid lines/group. Symbols in box plots represent the average of duplicate measures of individual colonoid lines. *=*P* < 0.05, **=*P* < 0.01, ***=*P* < 0.001.

Consistent with the metabolic profile of inactive UC colonoids and controls, only two genes (*HLA-DRA* and *HLA-DMB*) were significantly upregulated in iUC colonoids compared to control colonoids (Supplementary Datasheet 1).

Chemokines recruit immune cells to the epithelium in UC, where they can incite damage. We observed enhanced gene expression of chemokine genes involved in the recruitment of macrophages (CCL2), T-cells (CXCL11), neutrophils, (CXCL1), and T-and B-cells (CCL28) in aUC colonoids compared to controls (Figure 2*C*). Correspondingly, CCL2, CXCL11, and CCL1 proteins were disproportionately secreted into the medium in aUC compared to controls (Figure 2*D-F*). We then determined whether metabolic uncoupling during differentiation was sufficient to induce chemokine secretion in control colonoids. Indeed, treatment of control colonoids with Fccp during differentiation significantly upregulated the gene and protein secretion of CXCL1, CXCL11, and CCL2 (Supplementary Figure 4*O* and *P*). Thus, molecular events that stimulate metabolic uncoupling during epithelial differentiation in UC may contribute to the chemoattraction of immune cells.

### Active UC colonoids exhibit enhanced glycolytic capacity

To begin to understand the cellular underpinnings of hypermetabolism in aUC colonoids, we analyzed our differentially expressed transcriptomic data using functional annotation analysis (Ingenuity Pathway Analysis [IPA])^28^ to predict the dominant molecular and cellular functional differences between aUC and control colonoids. IPA implicated carbohydrate metabolism as the top mediator of cellular function during differentiation (Figure 3*A*, Supplementary Datasheet 2). Measurement of real-time extracellular acidification rates (ECAR)^29^ showed that aUC colonoids had an enhanced glycolytic capacity and reserve compared to control colonoids (Figure 3*B*-*E*). Still, non-glycolytic acidification in aUC was significantly higher compared to controls, (Figure 3*F*), which suggests other pathways such as CO_2_ from the TCA cycle contribute to extracellular acidification.^29^ To complement the extracellular flux analysis results, observed that on average, aUC colonoids consumed 4.2 times more glucose from the media and produced 2.8 times more lactate compared to controls over a 2-day differentiation (Figure 3*G* and *H*). Collectively, the data indicate that, during the early stages of differentiation, aUC colonoids possess enhanced glycolytic capacity which could support the observed hypermetabolism.

**Figure 3.**
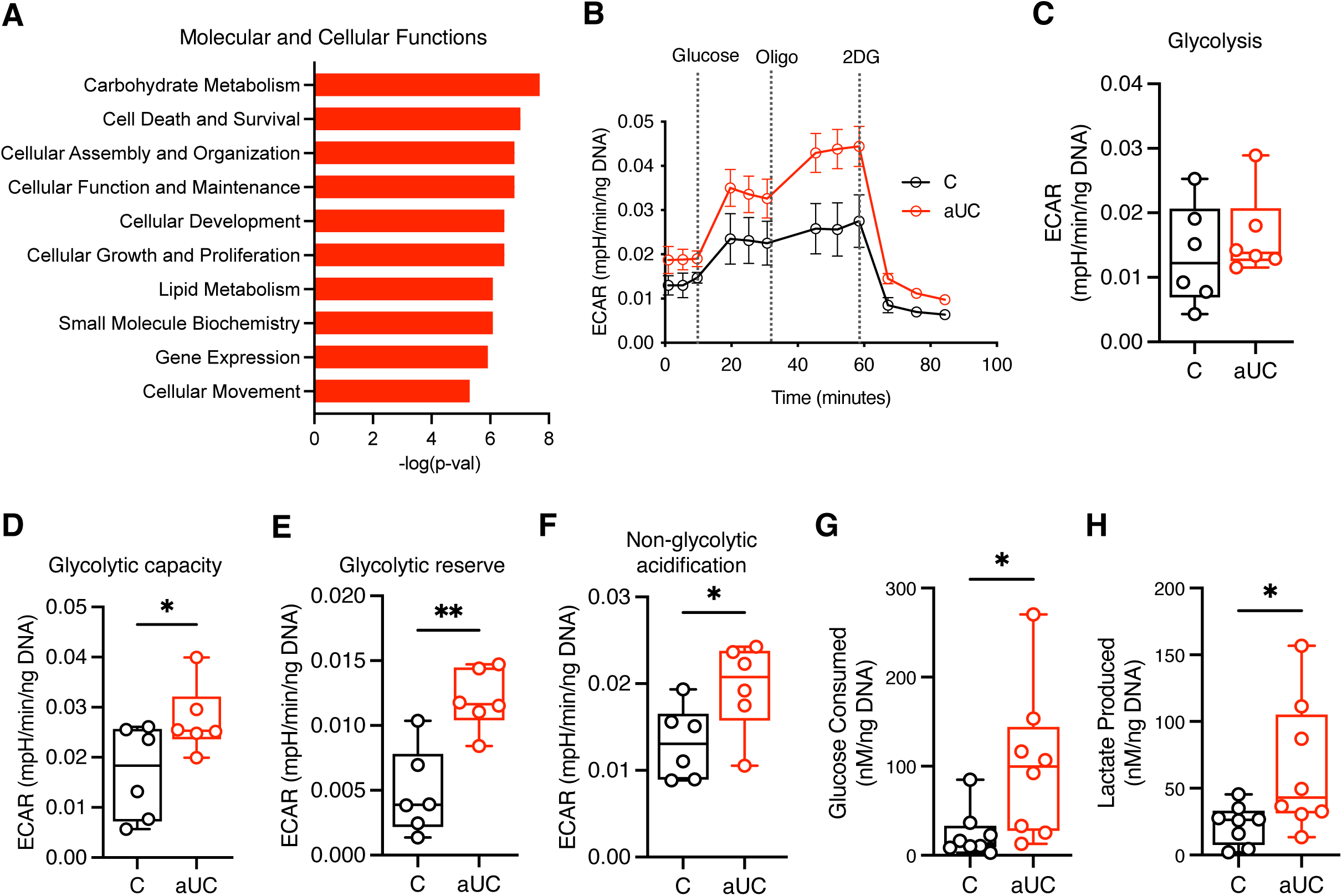
Higher glycolytic capacity in active UC colonoids during differentiation. (**A**) Ingenuity Pathway Analysis of differentially expressed RNAs between aUC and C colonoids. The top 10 Molecular and Cellular Functions are shown. Full details of molecules involved under each function are presented in Supplementary Datasheet 2. (**B**) Bioenergetic profile of C and aUC colonoids differentiated in 96-well Seahorse plates for 3 days and subjected to the Seahorse Glycostress assay. **(C-F**) Glycostress extracellular acidification rates (ECAR) of C and aUC colonoids as in (**B**) to determine real-time (**C**) glycolysis, (**D**) glycolytic capacity, (**E**) glycolytic reserve, and (**F**) non-glycolytic acidification rate. (**G**) Extracellular glucose and (**H**) extracellular lactate concentration in the medium from C and aUC colonoids differentiated for 2 days. Symbols in (**C-H**) represent the average measures of individual colonoid lines, n=6-8 patient colonoid line/group. *=*P* < 0.05, **=*P* < 0.01. ECAR, extracellular acidification rates.

### Dysregulated lipid metabolic genes in aUC colonoids

Functional annotation analysis of aUC colonoid differential gene expression also identified lipid metabolism as one of the top 10 dominant molecular and cellular functions (Figure 3*A*). Upregulated lipid metabolism genes in aUC included those related to fatty acid transport (*FABP6, ABCA12*), apolipoproteins (*APOL1, APOL3*), sphingolipids and phospholipid biosynthesis (*SPTLC3* and *DGKG*, respectively), lipid-metabolizing CYPs (*CYPC218*, *CYP3A4*), and lipolysis (*LPL, PLB1, AADAC, LIPA*). We examined these genes in other colon bulk and single-cell RNA-seq data from UC patients. First, the PROTECT bulk RNA-seq data^14^ showed enhanced expression of *ABCA12, APOL3, APOL1, LPL, DGKG,* and *LIPA* in active UC patients compared to controls (Supplementary Figure 5*A*). Next, we analyzed publically available UC single-cell RNA-seq data^30^ to evaluate whether inflamed epithelial cells exhibit similar upregulation of lipid-related genes in UC. Compared to healthy samples, a higher proportion of the inflamed samples visibly expressed these identified lipid-related genes including stem cells and other progenitor epithelial lineages (Supplementary Figure 5*B* and *C*). Taken together, these analyses of the organoid, bulk tissue, and single-cell RNA-seq data sets indicate the transcriptomic dysregulation of lipid-related genes in UC which is modeled in our colonoids.

### Aberrant lipid accumulation and oxidation in aUC colonoids during differentiation

Considering the altered expression of lipid metabolic genes in aUC, we sought to determine whether enhanced lipid metabolism contributes to the observed hypermetabolic phenotype. Consistent with their analogous metabolic states before differentiation, control, and aUC spheroids had similar neutral lipid content (Figure 4*B*). However, we observed a significant increase in neutral lipid accumulation during differentiation in aUC colonoids compared to their paired spheroids, while control spheroids and colonoids still retained similar neutral lipid content. (Figure 4*B*). Furthermore, differentiated aUC colonoids contained elevated lipid content compared to the control colonoids (Figure 4*B* and *C*). Our bulk RNA-seq data showed an upregulation of 4 lipid hydrolysis genes (*LPL, PLB1, AADAC, LIPA*) in aUC colonoids (Figure 4*A*) suggesting that hydrolysis of accumulated lipids to free fatty acids may undergo fatty acid oxidation (FAO) and support OXPHOS. We tested the protein activity of one of the lipid hydrolysis genes– LPL, and found elevated LPL activity in aUC colonoids compared to control on Day 2 with no significant difference between groups by Day 5 (Figure 4*D*). These data further suggest that active lipid cycling may be overamplified during differentiation in the UC colon epithelium and our pediatric colonoid model of UC may recapitulate these lipid metabolic dynamics.

**Figure 4.**
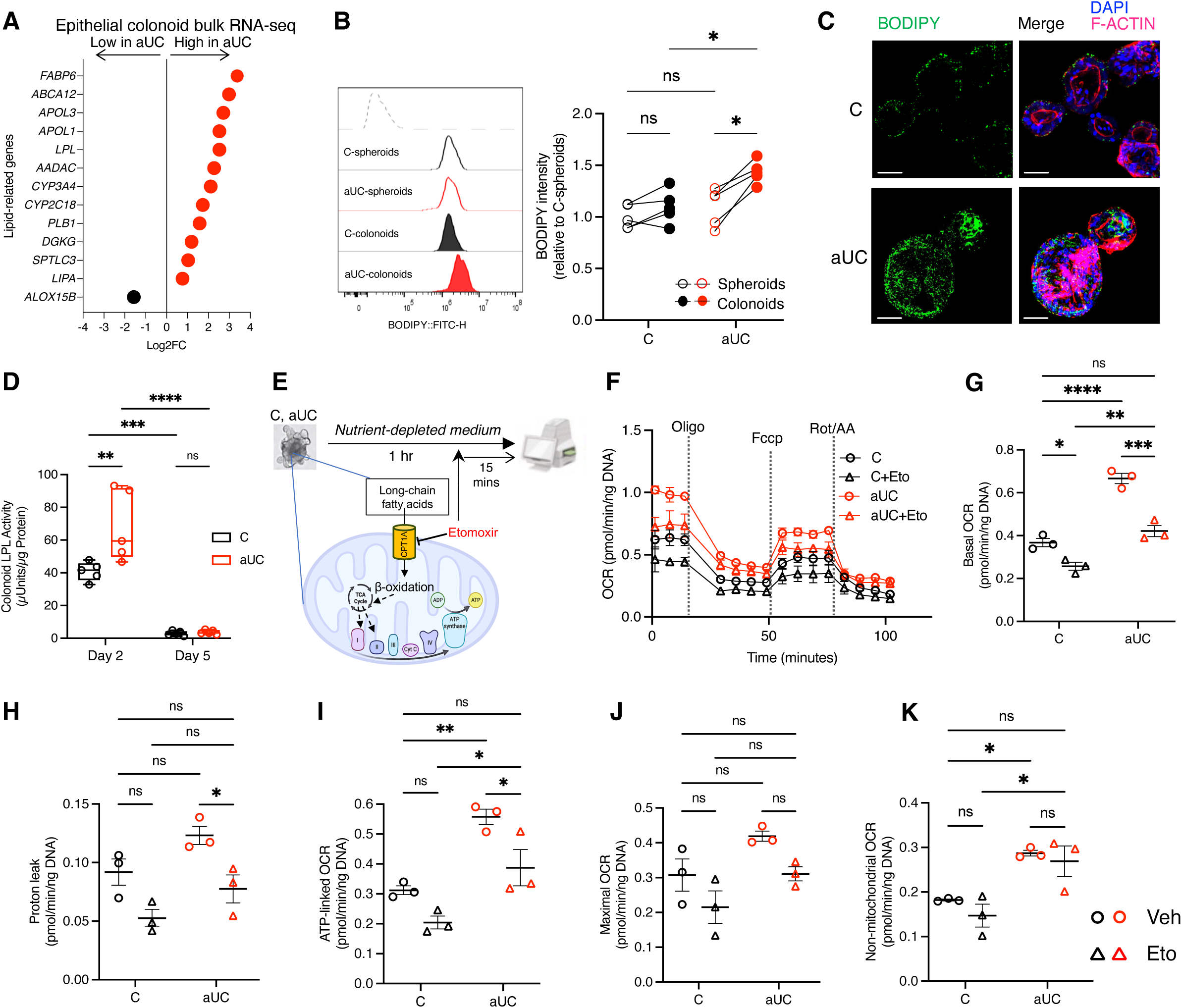
Dysregulated lipid metabolism supports hypermetabolism in active UC colonoids during differentiation. (**A**) Differentially expressed lipid-related genes (FDR<0.1) in aUC vs C epithelial colonoids. Dots represent the Log2 fold change in aUC compared to C colonoids. (**B**) BODIPY^+^ neutral lipid estimation by flow cytometry in spheroids (undifferentiated) and colonoids (differentiated). (**C**) Representative confocal microscopy images of C and aUC colonoids treated as in **B**, for visualizing BODIPY^+^ neutral lipid accumulation. Scale bar, 50 µm. (**D**) LPL protein activity in C and aUC colonoids after a 2-day and 5-day differentiation. (**E**) Schematic of C and aUC colonoids differentiated for 3 days and subjected to the Seahorse MitoStress with or without etomoxir. (**F**) Bioenergetic profile of C and aUC colonoids as in **E**. (**G**) Basal OCR, (**H**) Proton leak, and (**I)** ATP-linked OCR, (**J**) Maximal OCR, and (**K**) Non-mitochondrial OCR response of C and aUC colonoids as in **E**. For (**B** – **K**). Symbols represent the average measures of individual colonoid lines, n=3-8 patient colonoid lines/group *=*P* < 0.05, **=*P* < 0.01, ***=*P* < 0.001. ****=*P* < 0.0001. Eto, etomoxir; OCR, oxygen consumption rates.

Given the increased potential for neutral lipid accumulation and lipid hydrolysis in aUC colonoids, we hypothesized that the oversupply of lipids contributes to elevated O_2_ consumption in aUC colonoids during differentiation. Thus, we exposed control and aUC colonoids to a substrate-depleted medium with or without etomoxir, a carnitine palmitoyl-transferase 1a (CPT1a) inhibitor that inhibits fatty acid oxidation (Figure 4*D*). With negligible external sources of metabolic substrates, aUC colonoid mean basal OCR was 78% greater than that of controls (Figure 4*F* and *G*). Etomoxir reduced basal OCR in aUC colonoids to the level of control colonoids (Figure 4*G*). Interestingly, etomoxir significantly reduced proton leak in aUC but not in control colonoids, suggesting that enhanced FAO drives proton leak in aUC (Figure 4*H*). Compared to control colonoids, we observed a significant increase in ATP-linked OCR in aUC colonoids under this nutrient-depleted condition, with a significant decrease in ATP-linked OCR in aUC with etomoxir inhibition (Figure 4*I*). Etomoxir did not impact maximal respiration and non-mitochondrial respiration (Figure 4*J* and *K*). These results indicate that there is enhanced lipid accumulation and oxidation in the active UC epithelium which is necessary for the hypermetabolic phenotype during differentiation.

### PPAR-alpha contributes to hypermetabolic stress during colonoid differentiation

To understand the regulatory networks that may explain the hypermetabolic nature of differentiating aUC colonoids, we conducted a Causal Network Analysis^28^ of our colonoid bulk transcriptomic data. Among other notable molecular networks, the PPARA-RXRA network was the most significant causal factor (Figure 5*A*, Supplementary Datasheet 3). Analyses of lineage-specific PPARA expression in the scRNA-seq data from treatment-naïve UC colon epithelium^30^ showed the upregulation of PPARA majorly in progenitor cells (stem, TA2, cycling TA, secretory TA) in the inflamed and non-inflamed colon epithelial regions compared to healthy samples (Supplementary Figure 6*A*). We did not see any differences in *PPARA* mRNA expression in our bulk RNA-seq data between aUC and control colonoids differentiated for 3 days; however, we found an enhanced PPAR-α activity in nuclear extracts from aUC colonoids compared to controls at the early stages (24 hr) of differentiation (Supplementary Figure 6*B*).

**Figure 5.**
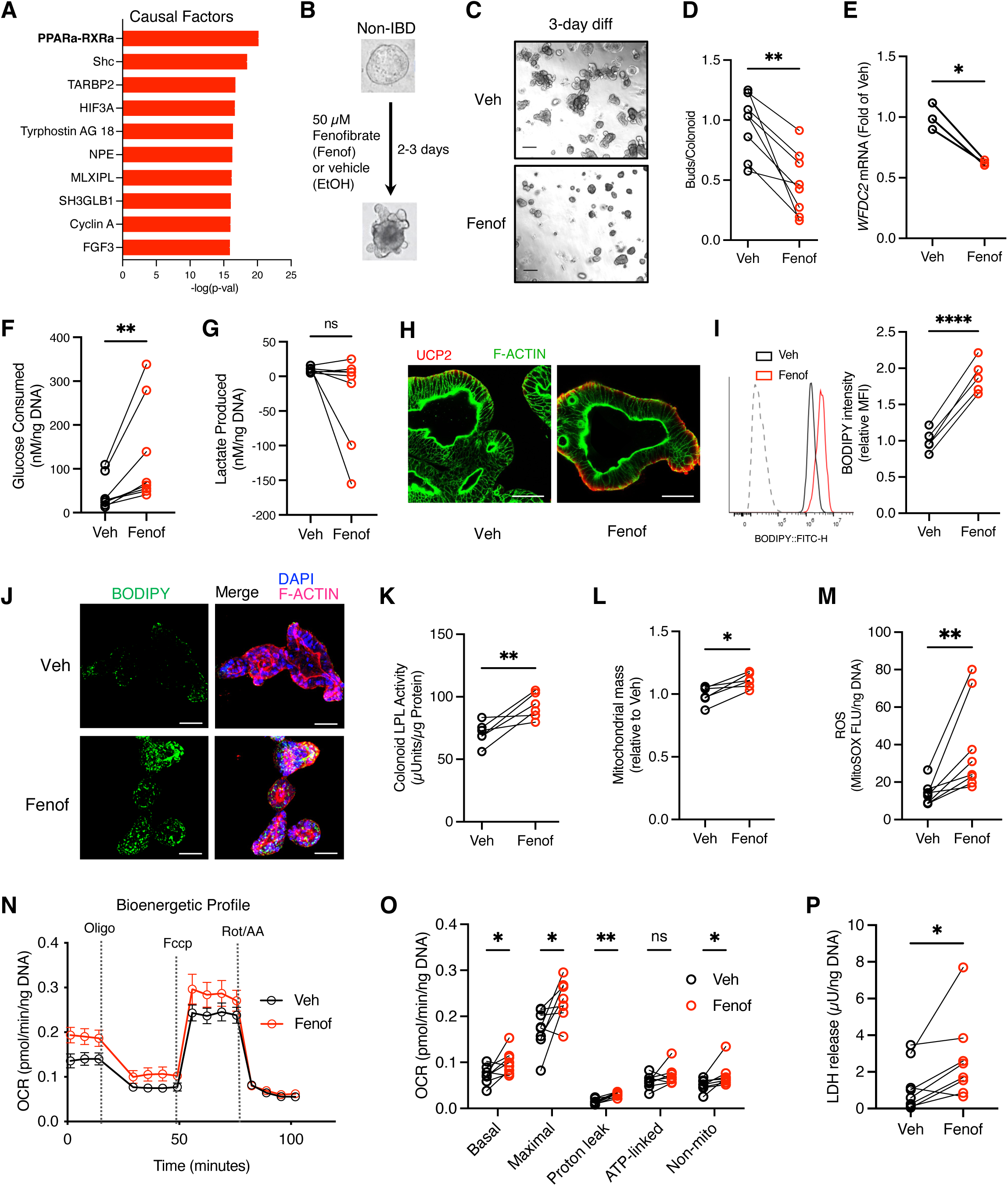
PPAR-α agonism in non-IBD colonoids induces hypermetabolic stress during differentiation. (**A**) Ingenuity Pathway Analysis of bulk transcriptomic data showing the top 10 predicted Causal Networks of differential gene expression between aUC and C colonoids. See Supplementary Datasheet 3 for all networks. (**B**) Schematic of PPARA agonist, fenofibrate, treatment in non-IBD colonoids during differentiation. (**C**) Representative phase contrast images of pediatric C colonoids treated as in **C**. Scale bar, 200 µm. (**D**) The number of buds/colonoid in colonoids treated as in **B**, and differentiated for 3 days. (**E**) *WFDC2* gene expression in colonoids treated as in **B**, assessed by qRT-PCR. (**F**) Extracellular glucose and (**G**) extracellular lactate concentration in medium from C colonoids treated with or without fenofibrate. (**H**) Representative confocal immunofluorescence images of UCP2 expression in non-IBD colonoids treated as in **B**, for 3 days. Scale bar, 50 µm (**I**) BODIPY^+^ neutral lipid estimation by flow cytometry. Data presented relative to the MFI of vehicle. (**J**) Representative confocal images of C colonoids visualizing BODIPY^+^ neutral lipids. Scale bar, 50 µm. (**K**) LPL activity in C colonoids treated as in **B**, and differentiated for 2 days. (**L**) Mitochondrial mass in C colonoids treated as in **B**, and differentiated for 3 days. (**M**) MitoSOX ROS estimate in C colonoids treated as in **B**, for 3 days. (**N**) Bioenergetic profile of C colonoids treated as in **B**, after a 3-day differentiation. (**O**) MitoStress OCR responses of C colonoids in **N**. (**P**) LDH activity in the medium during differentiation. For all data, symbols represent the average measures of individual colonoid lines, n=3-8 patient colonoid lines/group. *=*P* < 0.05, **=*P* < 0.01, ***=*P* < 0.001. ****=*P* < 0.0001. LDH, lactate dehydrogenase; OCR, oxygen consumption rates.

We wondered whether PPAR-α activation is sufficient to induce the observed hypermetabolic phenotype. Treatment of control colonoids with the PPAR-α agonist– fenofibrate during differentiation significantly reduced colonoid budded morphology and downregulated the crypt-base goblet cell marker^24^ *WFDC2* (Figure 5*B*-*E*). Fenofibrate enhanced glucose consumption vs vehicle which was not accompanied by significant lactate production, suggesting a negligible induction of anaerobic glycolysis (Figure 5*F* and *G*). Fenofibrate treatment enhanced UCP2 expression during colonoid differentiation (Figure 5*H*). Consistent with aUC lipid metabolic phenotypes, neutral lipid droplets, and LPL activity increased in control colonoids exposed to fenofibrate (Figure 5*I-K*). Fenofibrate enhanced mitochondrial mass, MitoSOX ROS, with a resultant elevation in basal, maximal, proton leak, and non-mitochondrial OCR with no significant increase in ATP-linked OCR (Figure 5*L*-*O*). Finally, fenofibrate treatment induced LDH release consistent with our observation in differentiating aUC colonoids (Figure 1*K*, Figure 5*P*). These findings suggest that overexpressed PPAR-α activity during colon epithelial differentiation significantly contributes to hypermetabolic stress.

### Pharmacological inhibition of PPAR-**α** reprograms epithelial metabolism in aUC colonoids

Following the observed contribution of PPAR-α activity to lipid accumulation and hypermetabolic stress during differentiation, we reasoned that inhibition of PPAR-α during aUC colonoid differentiation may ameliorate hypermetabolic stress. We treated aUC colonoids with the PPAR-α antagonist, GW6471 (GW) during differentiation. GW treatment suppressed neutral lipid accumulation and downregulated *LPL* and *UCP2* genes compared to untreated aUC colonoids (Figure 6*A*-*C*, Supplementary Figure 7*A*). Consequently, in a substrate-enriched medium (10 mM glucose, 1 mM pyruvate, 2 mM glutamine), we observed a significant reduction in basal, maximal, proton leak, and ATP-linked OCR in GW-treated colonoids (Figure 6*E*). Luminescence assay for total cellular ATP showed no significant differences in untreated and GW-treated aUC colonoids (Figure 6*F*). These data suggest that PPAR-α inhibition reduced lipid accumulation and improved cellular bioenergetics in active UC colon epithelial cells. Under substrate-depleted conditions, blockade of FAO with etomoxir did not affect cellular bioenergetics measures in GW-treated aUC colonoids (Figure 6*G* and *H*, Supplementary Figure 7*B-E*) which confirmed that GW abrogates the contribution of lipid accumulation and oxidation to cellular bioenergetics in aUC colonoids. Consequently, we observed a significant increase in glucose consumption in all GW-treated aUC colonoid lines compared to untreated aUC colonoids without changes in lactate production, suggesting an increased reliance on glucose as a substrate to drive cellular metabolism (Figure 6*I* and *J*).

**Figure 6.**
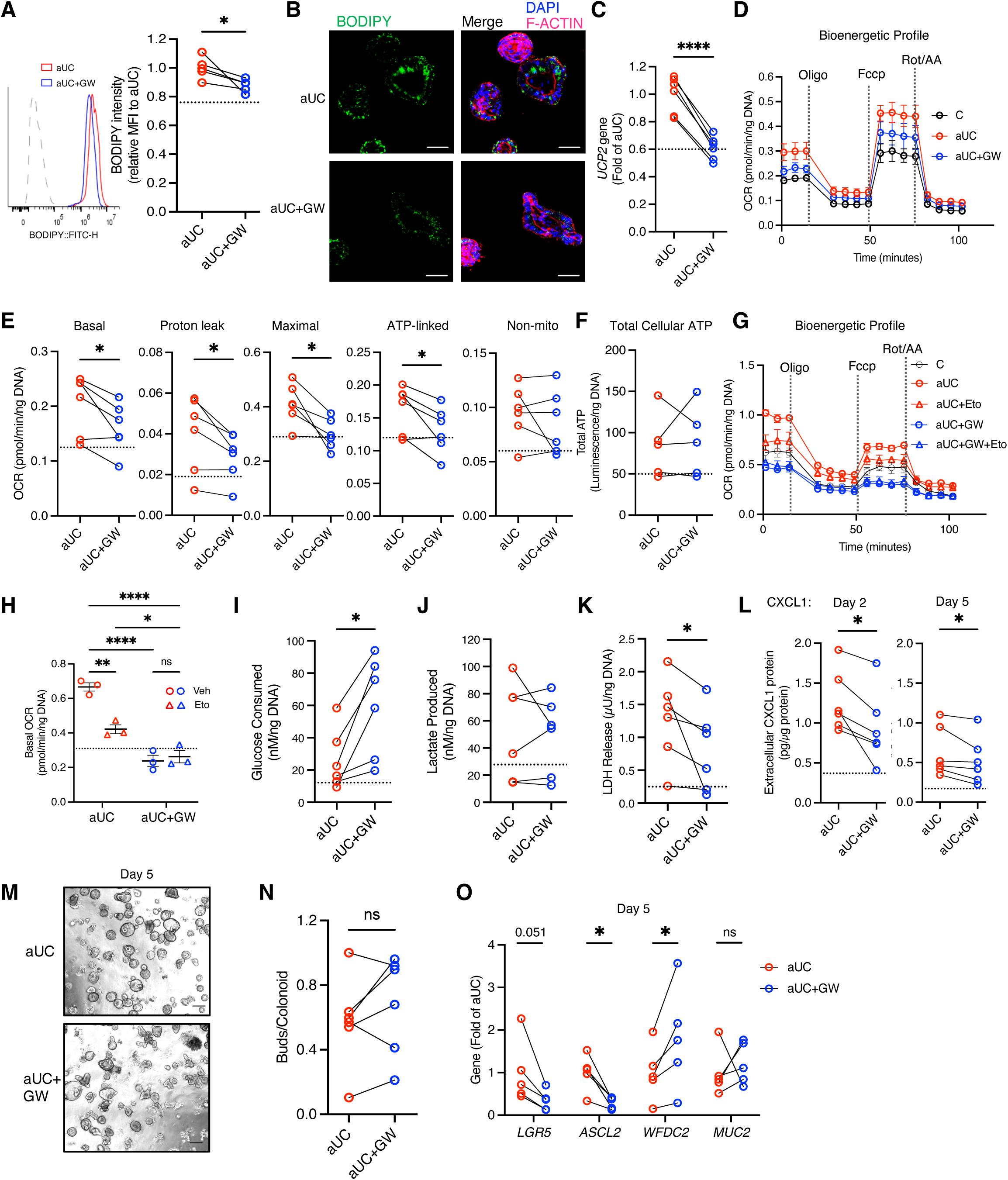
PPAR-α inhibition reprograms epithelial metabolism and ameliorates hypermetabolic stress in active UC colonoids during differentiation. aUC colonoids were treated with 1 µM of the PPAR-α antagonist, GW6471 (GW), or vehicle (EtOH) during differentiation. (**A**) BODIPY^+^ neutral lipid estimation in aUC colonoids. (**B**) Representative confocal images of aUC colonoids treated as in **A**, for visualizing BODIPY^+^ neutral lipid accumulation. Scale bar, 50 µm. (**C**) UCP2 gene expression by qRT-PCR. (**D**) Bioenergetic profile of aUC colonoids treated with or without GW and subjected to the Seahorse MitoStress test. (**E**) MitoStress OCR responses of aUC colonoids treated as in **D**. (**F**) ATP luminescence assay in colonoids cultured as in **D**. (**G**) Bioenergetic profile of aUC colonoids subjected to the Seahorse MitoStress test in a nutrient-deprived medium with or without etomoxir. (**H**) Basal OCR of aUC colonoids as in **G**. (**I**) Extracellular glucose and (**J**) lactate in the medium after 2 days. (**K**) LDH activity after a 2-day differentiation sampled from the same plate as **I**. (**L)** Medium collected on day 2 and on day 5 (representing the last 3 days of differentiation) was used to assess CXCL1 secretion. (**M**) Representative phase contrast images of aUC colonoids differentiated for 5 days. Scale bar, 200 µm. (**N**) The number of buds/colonoids in aUC colonoids treated as in **M**. (**O**) Gene expression of differentiation markers assessed by qRT-PCR. For all data, symbols represent the average measures of individual donor colonoid lines and dotted vertical lines represent the average of at least 2 non-IBD control colonoids, n=3-6 patient colonoid lines/group. *=*P* < 0.05, **=*P* < 0.01, ***=*P* < 0.001. ****=*P* < 0.0001 OCR, oxygen consumption rates; Eto, etomoxir

We next examined whether the metabolic reprogramming in aUC colonoids with PPAR-α inhibition impacted cellular health markers. GW treatment in aUC colonoids did not impact mitochondrial mass or mitochondrial ROS levels compared to untreated aUC colonoids (Supplementary Figure 7*F* and *G*). In contrast, GW treatment significantly reduced LDH release (cytotoxicity marker) in every colonoid line compared to untreated aUC colonoids (Figure 6*K*). In the medium collected from aUC colonoids differentiated for 2 or 5 days, we found that GW treatment significantly suppressed extracellular CXCL1 (Figure 6*L*). There was no significant effect of GW treatment on extracellular CXCL11 and a trend toward a reduction in extracellular CCL2 on day 2 (p=0.053) while there was no effect on day 5 (Supplementary Figure 7*H* and *I*). Blinded visual assessment of colonoid morphology showed no consistent impact of GW on colonoid budding (Figure 6*M* and *N*). The gene expression of stemness markers (*LGR5* and *ASCL2*) reduced over time in every aUC colonoid line with GW treatment, with significant downregulation in *ASCL2* (Figure 6*O*). *WFDC2* significantly increased in GW-treated colonoid lines compared to untreated aUC colonoids while the increase in *MUC2* expression was not statistically significant (Figure 6*O*). Overall, these data suggest that PPAR-α inhibition in aUC epithelial colonoids induced a metabolic shift away from lipid oxidation which consequently reduced chemokine secretion, and augmented molecular markers of epithelial differentiation.

## DISCUSSION

Leveraging clinical samples to develop human IBD pre-clinical models is important for determining mechanisms of human disease pathogenesis. We used patient-derived colon epithelial organoids as a human model to study epithelial metabolic dysfunction in pediatric UC. We report profound elevation in oxygen consumption in aUC colonoids during differentiation related primarily to increased proton leak. This hypermetabolic phenotype was associated with increased chemokine secretion, oxidative stress, cytotoxicity, and impaired differentiation. The aUC colonoids showed an elevated capacity for lipid accumulation during differentiation and their hypermetabolic phenotype required FAO. Our data indicate PPAR-α drives altered lipid metabolism and OXPHOS in aUC colonoids and that pharmacological inhibition of PPAR-α normalizes abnormal metabolism, chemokine expression, and differentiation in aUC epithelial cells.

In our study, colonoids from active UC patients showed increased mitochondrial respiration during differentiation without a concomitant change in ATP levels. The normal colon epithelium is less dependent on highly oxidative processes for energy generation.^31,32^ Accordingly, our data in non-IBD control colonoids showed marginal increases in median oxygen consumption during the energy-demanding differentiation process. Hypermetabolism may channel energetic demands towards molecular programs that favor a hypersecretory phenotype,^33,34^ which may explain the hypersecretion of chemokines in aUC colonoids during differentiation. Furthermore, the oversupply of metabolic substrates may result in hypermetabolism with associated mitochondrial and cellular stress.^35^ Cellular dependency on lipids for energy generation is inefficient since the oxidation of lipids consumes energy and induces metabolic uncoupling^36^ as observed with increased proton leak in our aUC colonoids. Conversely, aberrant lipid accumulation or lipid droplet hydrolysis may trigger lipotoxicity-related pathways including apoptosis, inflammation, and ER stress.^37^ Overall, our data suggest a lipid-related hypermetabolism during differentiation in epithelial aUC colonoids, which may increase the energetic cost of differentiation and contribute to enhanced epithelial cellular stress in aUC. Hence, it is plausible that approaches that suppress aberrant epithelial lipid accumulation, hydrolysis, and oxidation may improve colon epithelial function in UC.

The large PROTECT pediatric UC inception cohort study^14^ found extensive downregulation of genes related to mitochondria and OXPHOS in the rectal mucosa. While we did not observe any significant changes in mitochondrial genes in our aUC colonoids, we observed abnormally high mitochondrial respiration in aUC colonoids. Similarly, enhanced epithelial OXPHOS activity was reported in a mouse model of colitis during the active stage of the disease at day 7 despite the downregulation of some mitochondrial genes.^38^ These metabolic and transcriptional disparities in the active epithelium largely resolved to the level of control upon recovery at day 28.^38^ There may be other contributors to epithelial metabolic dysregulation in UC, including the colon mucosa’s inflammatory milieu, and it may be that our colon organoids predominantly retained the lipid-dependent components in culture.

Our study also suggests a role for the PPAR-α network in the UC epithelium and colonoids. PPAR-α activation in non-IBD colonoids during differentiation recapitulated lipid-induced hypermetabolic stress, morphologic and molecular evidence of impaired differentiation, similar to our observation in aUC colonoids. Consistent with our study, PPAR-α agonism exaggerated colitis in DSS-induced mouse models with a concomitant increase in bioactive serum lipids.^39,40^ Furthermore, activation of fatty acid oxidation partly via PPAR-α promotes intestinal stemness in high-fat diet models,^41^ suggesting a detrimental effect on intestinal stem cell differentiation and maintenance of epithelial homeostasis. Natural PPAR-α agonists including PUFAs are disproportionally accumulated in the inflamed colon mucosa of UC patients which correlated positively with endoscopic disease activity.^42,43^ In contrast, pharmacological inhibition of PPAR-α in our aUC colonoids suppressed aberrant neutral lipid accumulation and oxidation. In this scenario, aUC colonoids consumed more glucose likely due to a metabolic switch to compensate for the loss of lipid-dependent oxidation for energy, resulting in reduced cytotoxicity and CXCL1 secretion. Hypermetabolism may divert energy expenditure towards translational processes that upregulate cytokine and stress responses;^33^ hence, it is plausible that the blockade of lipid’s contribution to the hypermetabolic state in aUC colonoids by PPAR-α inhibition reduces such cytotoxic stress responses. Animal studies have suggested the potential of PPAR-α inhibition in ameliorating colitis.^44,45^ Overall, our study in human patient-derived colonoids suggests that PPAR-α antagonism in the epithelium may be a viable treatment strategy for reducing the contribution of lipotoxicity to epithelial stress in UC.

In conclusion, our data in patient-derived epithelial colon organoids indicate that hypermetabolism and lipid metabolic dysregulation significantly contribute to colon epithelial dysregulation in pediatric UC. Furthermore, epithelial PPAR-α may be a therapeutic target for restoring colon epithelial homeostasis by reducing dysregulated lipid metabolism and normalizing cellular respiration. The use of patient-derived colonoids in this study opens novel research directions in colon epithelial metabolism that may identify metabolic targets for next-generation therapies specifically directed toward healing the colon epithelia in pediatric UC.

## Supplementary Materials

Supplementary Materials and Methods

Supplementary Figures 1-7

Supplementary Table 1 and 2 Table S1 to S2

Supplementary References

Supplementary Datasheets 1 to 3 (Excel files)

## Supporting information

Supplementary Datasheet 1

Supplementary Datasheet 2

Supplementary Datasheet 3

## Acknowledgments

We thank L. Prince (Pediatric Neonatology, Stanford Medicine) for resources for flow cytometry and confocal microscopy. We acknowledge the support of the National Center for Advancing Translational Sciences of the NIH under Award Number UL1TR003142, and support by the Stanford Innovative Medicines Accelerator (IMA-1494). Some illustrations were prepared with biorender.com.

## Author contributions

Conceptualization: B.A.O., S.D., and M.J.R.

Organoid biobank generation: B.A.O., S.R.F., A.W., M.G., T.T., A.L.D., and M.J.R.

Methodology: B.A.O., S.R.F., M.E.M., S.D., V.D.J.P., and M.J.R

Investigation: B.A.O., L.H., M.G., E.I.A.E., and N.S.

Analysis and Visualization: B.A.O., L.H., and J.A.L.

Project administration: B.A.O., A.L.D., V.D.J.P., and M.J.R

Supervision: M.J.R.

Writing – original draft: B.A.O.

Writing – review & editing: B.A.O., M.J.R.

## Manuscript approval: All authors

Funding acquisition: S.D., A.L.D., and M.J.R.

## Data Availability

Bulk RNAseq data sets from spheroids and colonoids in this study were deposited into Gene Expression Omnibus (GEO) under the accession number GSE276170. Other appropriate data will be provided by the corresponding author upon request.

