## Supplementary Methods and Figures for "Patient-derived colon epithelial organoids reveal lipid-related metabolic dysfunction in pediatric ulcerative colitis"

Babajide A. Ojo et al.

**This file includes:**

Supplementary Materials and Methods

Supplementary Figures 1-7

Supplementary Table 1 and 2 Table S1 to S2

Supplementary References

**Other Supplementary Material for this Manuscript includes the following:**

Supplementary Datasheets 1 to 3 (Excel files)

**SUPPLEMENTARY MATERIALS AND METHODS**

**Study Design and Clinical Samples**

Pediatric patient recruitment occurred at Cincinnati Children’s Hospital Medical Center, Cincinnati, Ohio, and Lucile Packard Children’s Hospital Stanford, Palo Alto, California. Patient data and biopsy collection followed approved protocols by the Research Ethics Committees at each institution. Biopsies from patients with active UC, inactive UC, and non-IBD controls were collected during clinically-indicated lower endoscopy and transported on ice to the lab for immediate processing for organoid generation. Morphological measures were conducted by an investigator blinded to the diagnosis. Patient demographics and clinical data are presented in Supplementary Table 1.

**Biopsy processing and organoid culture**

Fresh biopsies were transferred into 5 mL of 2 mM PBS-EDTA and incubated for 45 min on a rotator in a cold room. Biopsies were transferred into 2.5 mL ice-cold sorbitol buffer (2% sorbitol, 1% sucrose, 1% BSA, 50 µg/mL Normocin) in a petri dish and crypts were gently scraped off using a college cotton tweezer under a dissecting microscope (Leica S9E). Big tissue pieces were removed with a 20 µL pipette tip and the crypts and buffer were transferred into a 15 mL tube. Crypts were sheared briskly with a 1000 µL pipette (80X total with 1 min interval) and the suspension was made up to 14 mL with ice-cold PBS (Gibco #10010-023). The suspension was centrifuged (300 x g, 5 mins, 4^o^C) and the pellet was resuspended in organoid wash medium, OWM, (DMEM/F12, 10% FBS, 50 µg/mL Normocin, and washed twice (300 x g, 5 mins, 4^o^C). The visible pellet was resuspended in growth factor-reduced Matrigel (Corning #356231) and 50 µL drops of the Matrigel mixture were allowed to polymerize at 37^o^C for at least 15 mins on pre-warmed 12-well plates. Upon stabilization, 700 µL of Intesticult organoid growth medium, OGM, (STEMCELL Technologies, #06010) containing 10 µM TGF-beta receptor inhibitor, SB431542 (Cayman Chemical, #13031), 10 µM ROCK inhibitor, Y27632 (STEMCELL Technologies, #72304), and 50 µg/mL Normocin. Crypts were placed in a tissue culture incubator (37^o^C, 5% CO2) and OGM was replaced every 2-3 days for the first 7-14 days. Spheroids were considered stable and mature enough for expansion when >70% of spheroids are ~100 µm in diameter.

For spheroid expansion, OGM was aspirated and 1mL of ice-cold PBS was added to each well. The Matrigel domes were dislodged from the bottom of the plate with a 1000 µL pipette and the PBS-Matrigel mixture was transferred into a 15 mL tube. The Matrigel was broken up inside the tube with a 1000 µL pipette, made up to 5 mL with ice-cold PBS, and incubated on ice for 5 min. The suspension was centrifuged (300 x g, 5 mins, 4^o^C), 350 uL of TrypLE Express (Gibco #12604-013) was added to the spheroid pellets and incubated in a 37 ^o^C water bath for 4-5 mins. Spheroids were broken up by pipetting 50X with a 1000 µL pipette followed by the addition of 5 mL ice-cold OWM. Spheroids were washed twice in OWM (300 x g, 5 mins, 4^o^C), the resulting pellets were resuspended in 3 parts of Matrigel to 1 part of OWM, and 50 µL drops of the Matrigel suspension were plated in pre-warmed 12-well plates and allowed to polymerize as previously described. For spheroid expansion, 2-3 drops of this suspension were plated per well of a 12-well plate and 800 µL of L-WRN conditioned medium^1,2^ was added and replaced every other day.

For cryopreservation, spheroids underwent at least 3 passages before storage in our pediatric organoid biobank. Matrigel domes with matured spheroids (~80 µm) were dislodged in ice-cold PBS and centrifuged as previously described. Pellets were resuspended in freezing medium (90% FBS, 10% DMSO), and 1 mL of the suspension was added to a cryotube. Samples were slow-frozen in a Mr. Frosty freezing container at -80^o^C and transferred to liquid N_2_ for long-term storage. Overall, the pooling of matured spheroids from two Matrigel domes in one cryotube allows for sufficient post-thaw recovery of spheroids.

All experiments used cryopreserved spheroids. Withdrawal of cryotubes from the LN_2_ tank followed a quick thaw in a 37^o^C water bath for ~2mins. Cryotube contents were quickly transferred into pre-warmed 6 mL of OWM and centrifuged (300 x g, 5 mins, 20^o^C). Spheroid pellets were immediately resuspended in 5 mL OWM, washed once, resuspended in a 3:1 mix of Matrigel and OWM, and plated in 50 µL domes as previously described. Cryopreserved spheroids were passaged at least once before usage for all analyses.

**Spheroids processing and plating for analysis**

After 5-7 days after passaging with TrypLE Express, Matrigel domes with matured spheroids were dislodged with ice-cold PBS as previously described and filtered with a 100 µm strainer (pluriSelect). Filtrates were pelleted and washed once in OWM, manually counted, and replated in domes containing a 1:1 mix of Matrigel and OWM (8-15 spheroids per µL depending on the analysis) and maintained in OGM. For all analyses requiring differentiation into colonoids, replated spheroids were allowed to acclimatize overnight in OGM before switching to Intesticult human organoid differentiation medium, ODM, (STEMCELL Technologies, #100-0214) for the indicated differentiation durations with ODM changed every 2 days. In some experiments, matured spheroids were filtered through a 40 µm strainer (Pluriselect) immediately after passaging and cultured in OGM for 6-7 days before counting, replating, and differentiation when necessary.

To measure drug-induced responses, 2 µM FCCP (Sigma, #C2920), 50 µM Fenofibrate (Abcam, AB120832), and 1 µM GW6471 (Sigma, #G5045) were added to ODM throughout the differentiation as indicated in figure legends. The solvents served as vehicle controls for respective experiments (DMSO for FCCP, and 100% ethanol for Fenofibrate and GW6471). ODM with drugs were removed and colonoids were processed for analyses.

**Spheroid Diameter and Colonoid Budding**

Spheroids from eight patients per group were passaged as previously described and filtered through a 40 µm strainer (Pluriselect). Spheroids were manually counted and 750 spheroids were replated per 50 µL dome in pre-warmed 12-well plates and cultured in OGM for 6 days. Phase contrast images were obtained on the Keyence BZ-X800 microscope.

To assess colonoid budding, matured spheroids were counted, replated in 50 µL Matrigel (15 spheroids/µL), and allowed to acclimatize overnight in OGM. Each well was washed with PBS at room temperature (RT) for 3 min before switching to ODM for the indicated number of days. Phase contrast images were obtained on the Keyence BZ-X800 microscope. All measurements of spheroid diameter and enumeration of colonoid budding were performed in a manner blinded to the colonoid diagnosis group using the Keyence BZ-X800 analyzer. The number of buds on each colonoid was presented relative to the total number of visible colonoids per field. The data presented represents an average of 20-30 colonoids per patient sample.

**Extracellular Flux Analyses**

The XF Seahorse MitoStress Test (Agilent) was used to measure oxygen consumption rates (OCR), at baseline, and after the injection of mitochondrial inhibitors.^3^ Each spheroid line was replated in quadruplicates as described previously in 15 µL matrigel domes (8 spheroids/µL) containing a 1:1 mix of Matrigel and OWM into Seahorse XF96 cell culture plates (Agilent #101085-004) and cultured overnight in 150 µL OGM. The four edges of the plate served as blank with only a 1:1 mix of Matrigel and OWM. For some experiments, 50 µL of OGM was removed after 24 hrs into an empty 96-well plate and stored at -80^o^C for later measurements of LDH activity. Where differentiation was needed, spheroids were washed once with PBS on the following day and differentiated in 200 µL ODM into colonoids for the indicated number of days in the figure legends.

On the day of the assay, ODM was removed from colonoids, washed once with 200 µL PBS, and replaced with 150 µL DMEM assay medium (Agilent, #103334-100), pH 7.4 containing 10 mM glucose, 2 mM glutamine, and 1 mM pyruvate (all from Agilent). Plates were incubated in a non-CO_2_ incubator for 1 hr and OCR was measured at baseline and after a sequential injection of oligomycin (final concentration, 4µM), FCCP (final concentration, 2 µM), and rotenone/antimycin (final concentration, 100 nM/1 µM) using the XF96^e^ Extracellular Flux assay kits on the XF96^e^ analyzer.

The Glycostress test (Agilent) was used to measure extracellular acidification rates (ECAR) in colonoids after a 3-day differentiation. On the day of the assay, ODM was replaced with base medium (Agilent #103575-100) lacking glucose, pyruvate, and glutamine and incubated in a non-CO_2_ incubator for 1 hr. Baseline ECAR was assessed using the XF96^e^ analyzer (Agilent) followed by sequential injections of glucose, oligomycin, and 2-deoxy glucose at final concentrations of 40 µM, 4 µM, and 500 mM, respectively. Raw data were normalized with DNA content as described below.

For etomoxir experiments, colonoids were differentiated for 3 days. On the day of the assay, ODM was replaced with assay medium (Agilent #103575) lacking glucose, pyruvate, and glutamine, and incubated in a non-CO_2_ incubator for 40 min. Etomoxir (Sigma, #E1905) was prepared in the assay medium, quickly added to designated wells within 5 mins at a final concentration of 44 µM, and returned to the non-CO_2_ incubator for 15 min. OCR assessment followed as described previously.

**DNA extraction**

For the normalization of colonoid extracellular flux data, DNA normalization protocol was used as previously described with modifications ^3^. DNA was extracted using the QIAamp DNA Micro kit (Qiagen #56304). Briefly, the medium was removed from the cell culture plates and 45 µL of ATL lysis buffer was added per well, followed by 5 µL of Protein kinase K. The plate was incubated at 56^o^C for 1 hr, quadruplicate wells were pooled into 1.5 mL tubes and returned to 56^o^C incubator for 1 hr. DNA extraction continued according to the manufacturer’s instructions. The DNA content of each colonoid line was calculated relative to the blank wells and used to normalize raw data.

**Total ATP measurement**

Spheroids were plated in duplicates in clear flat bottom 96-well tissue culture plates (Falcon #353075) as described for extracellular flux analyses and differentiated for the indicated number of days. Total ATP levels were assayed with the luminescent ATP detection assay kit (Abcam, ab113849), according to the manufacturer’s instructions and luminescence was recorded using the Spectramax id5 plate reader (Molecular Devices). ATP levels were normalized by measuring DNA content in a parallel plate as described for extracellular flux analyses.

**Mitochondrial Superoxide (MitoSOX) Analysis**

Spheroids were plated in duplicates in black 96-well tissue culture plates with clear bottoms (Corning, #3603) as described for extracellular flux analyses and differentiated for the indicated number of days in the figure legends. Mitochondrial ROS was estimated with the fluorescent probe MitoSOX red (Thermofisher, #M36008). Briefly, a 5 mM MitoSOX stock solution was prepared in DMSO and diluted with HBSS to a working concentration of 3 µM. Colonoids were washed twice with HBSS and the MitoSOX working solution was added to the colonoids and incubated for 30 min (37^o^C, 5% CO2). The colonoids were washed 3 times with HBSS and MitoSOX fluorescence was measured at 396/610 nm using the Spectramax iD5 plate reader (Molecular Devices). Data were normalized by measuring DNA content as described for extracellular flux analyses.

**Lactate Dehydrogenase (LDH) release**

Spheroids were plated in quadruplicates in 96-well plates as described for extracellular flux analyses and differentiated in ODM. After 48 hrs, 100 µL of ODM was collected in an empty 96-well plated and stored frozen at -80^o^C for later measurements of LDH activity. Extracellular LDH activity was measured using the LDH-Glo Cytotoxicity assay (Promega, #J2380). Samples were diluted 40X in ice-cold storage buffer (200mM Tris-HCl (pH 7.3), 10% Glycerol, 1% BSA), and LDH activity was measured according to the manufacturer’s instructions. Luminescence was recorded using the Spectramax id5 plate reader (Molecular Devices), and the data was normalized with DNA content from the same plate as described previously. LDH release from spheroids and colonoids was estimated relative to the medium in Matrigel-only blank wells.

**Extracellular Glucose and Lactate**

ODM from colonoids stored from 96 cell culture plates as previously described for LDH activity were thawed on ice and extracellular glucose and lactate were measured using the Glucose-Glo (Promega, #J6021) and Lactate-Glo (Promega, #J5021) assays, respectively. Samples were diluted 50X in ice-cold PBS and the measurements continued according to the manufacturer’s instructions. Luminescence was recorded using the Spectramax id5 plate reader (Molecular Devices), and the data was normalized with DNA content from the same plate as previously described. Glucose consumed or lactate produced were estimated relative to the medium in Matrigel-only blank wells.

**Flow Cytometry**

Mitochondrial mass (mtMass) analyses

For mtMass analyses, we labeled the mitochondria in spheroids and colonoids using the MitoTracker Green FM probe (Cell Signaling, #9074). Briefly, single cells from matured spheroids (15 spheroids/µL) and corresponding colonoids were obtained by the addition of 900 µL TrypLE Express followed by incubation at 37^o^C for 10 mins. Single cells were released by vortexing for 20 seconds with 10-second intervals, and washed in FACS buffer (5% FBS in PBS). Pellets were resuspended in 150 nM mtGreen solution prepared in phenol-red free DMEM/F12 + 5% FBS, followed by incubation for 30 mins at 37^o^C in a 5% CO_2_ incubator. Cells were washed twice in 2 mL PBS and resuspended in 500 µL of ice-cold PBS. The cells were passed through Falcon FACS tubes with a cell strainer cap (Fisher, #0877123) and incubated with 5 µL of the live-dead stain, 7-Aminoactinomycin D (7-AAD), for 5 mins on ice. The mean fluorescence intensity (MFI) of mtGreen was measured in 5000 events using the FITC channel and following 7-AAD exclusion (PE-Cy5) on the Agilent NovoCyte Flow Cytometer. Analysis and images were obtained using the FlowJo software v.10

Mitochondrial Membrane Potential (MMP)

Single cells from colonoids were processed as previously described and MMP was assessed with the MitoProbe JC-1 assay kit (Thermo Fisher Scientific, MP34152) following the manufacturer’s protocol. Briefly, Cells were incubated with 1 µM JC-1 in PBS for 30 mins at 37^o^C, 5% CO_2_. The cells were washed once with PBS and passed through Falcon FACS tubes with a cell strainer cap and incubated with 5 µL of 7-AAD, for 5 mins on ice. 10000 events from each line were recorded using the Agilent NovoCyte Flow Cytometer. After 7-AAD exclusion, MMP was calculated as the MFI ratio of the red (PE)/green (FITC) channels.

BODIPY Neutral lipids

Matured spheroids from each line were replated (15 spheroids/µL) in two wells of a 24-well plate and allowed to stabilize overnight in OGM. For each line, one well was switched to ODM to initiate differentiation into colonoids for 2 days while the other well continued in OGM. Single cells were processed from spheroids and colonoids from each line as previously described and cell pellets were fixed using the Cytofix/Cytoperm kit (BD Biosciences, #555028) according to the manufacturer’s instructions. Neutral lipid droplets in cells were stained with 1µg/mL BODIPY 493/503 (Thermo Fisher Scientific, #D3922) in PBS for 1 hr at RT with gentle mixing every 20 mins. Cells were washed once with 2 mL PBS and passed through Falcon FACS tubes with a cell strainer cap. BODIPY MFI was recorded in 5000 events using the FITC channel on the Agilent NovoCyte Flow Cytometer. Analysis and images were obtained using the FlowJo software v.10.

**Immunofluorescence Imaging**

Colonoid Embedding in Optical Cutting Temperature (OCT) compound

Matured spheroids from each line were replated (15 spheroids/µL, 3:1 mixture of Matrigel and OWM) as previously described in 6 domes per well of a 6-well plate. After a 3-day differentiation, colonoids were fixed with 5 mL of 2% Paraformaldehyde and 0.1% Glutaraldehyde for 30 mins at RT. Colonoids were washed 3X with 5 mL PBS for 10 min, and each Matrigel dome was carefully scooped into a 50 mL tube with 20% sucrose in PBS. After 2-3 days following the sinking of all domes, 6 domes/line was embedded in Tissue Tek OCT compound (Sakura Finetek, #4583) and frozen in liquid N_2_ using the Seal’N Freeze cryotray and box (Fisher Scientific, #NC1877501). From the Colonoid OCT block, 10 µm thick cryosections were cut on slides and stored at -80^o^C. For immunostaining, cryosections were washed once with PBS for 15 secs at RT followed by autofluorescence quenching by incubation with 10 mM Sodium Borohydride in PBS twice for 5 mins at RT. Sections were washed thrice in PBS for 10 minutes each, followed by permeabilization with 0.15% Triton X-100 for 15 mins at RT. Slides were washed thrice with PBS and incubated with the blocking buffer (3% BSA, 5% donkey serum, and 5% goat serum in PBS) for 1 hr at RT. This was followed by overnight incubation with primary antibodies rabbit anti-UCP2 (1:100, Proteintech, #11081-1-AP) and mouse anti-COX4 (1:200, Invitrogen, GT6310) in the blocking buffer in a humidified chamber at 4^o^C. Slides were washed twice with PBS for 15 secs, followed by 2X wash in PBS for 5 mins. Slides were incubated in a humidified chamber for 1 hr at RT in secondary antibodies: donkey anti-rabbit Alexa Flour 647 (1:1000, Invitrogen, #A31573) and goat anti-mouse Alexa Flour 488 (1:1000, Invitrogen, #A11001) prepared in blocking buffer. Slides were washed 3X in PBS and counterstained with Phalloidin (1:250, ThermoFisher Scientific #T7471) in PBS for 30 min at RT. Slides were washed once in distilled water and mounted in Prolong Glass Antifade Mountant with NucBlue Stain containing the Hoechst 33342 DNA marker (ThermoFisher Scientific, #P36983). Images were acquired on the Keyence BZ-X800 microscope. Images were merged with the Keyence BZ-X800 software analyzer and cells expressing DAPI and those co-expressing UCP2 and COX4 were counted on at least four high-power fields (60X oil objective) per patient colonoid line.

Colonoid whole-mount staining

For whole-mount staining, matured spheroids were replated (15 spheroids/µL, 1:1 mixture of Matrigel and OWM) as previously described in 8-well chamber µ-slides (IBIDI, #80826). After differentiation, colonoids were washed with PBS at RT and fixed with 500 µL of 4% PFA for 30 mins at RT. For UCP2 staining, colonoids were washed 2X with 500 µL PBS for 5 mins and permeabilized with 0.2% Triton X-100 in PBS for 30 mins at RT followed by 2X wash in PBS. Samples were incubated with 50 mM ammonium chloride (NH_4_Cl) in PBS for 30 mins at RT to quench potential autofluorescence. After a 2X wash in PBS, samples were incubated in blocking buffer (3% BSA, 5% donkey serum in PBS) for 1 hr at RT and incubated in primary antibody- rabbit anti-UCP2 (1:50, Proteintech, #11081-1-AP) overnight and secondary antibody- donkey anti-rabbit Alexa Flour 647 (1:500, Invitrogen, #A31573) for 1 hr at RT, and incubated with Phalloidin as previously described. Samples were mounted in 200 µL Fructose-glycerol clearing solution ^4^ and incubated for 30 mins at RT. Whole-mount images were acquired using the Stellaris 8 Inverted Confocal Microscope (Leica) using the 20x oil objective and individual channels were merged using the Leica LAS X software.

For visualizing neutral lipids, colonoids differentiated for 2 days in chamber slides were fixed in 4% PFA, washed, and incubated in 50 mM NH_4_Cl as previously described. Then colonoids were stained with 1 µg/mL BODIPY 493/503 (Thermo Fisher Scientific, #D3922) in PBS for 1 hr at RT, washed twice in PBS, and incubated with Phalloidin as previously described. Chamber slides were washed once in distilled water and mounted in 2 drops of Prolong Glass Antifade Mountant with NucBlue Stain containing the Hoechst 33342 DNA marker (ThermoFisher Scientific, #P36983). Samples were allowed to cure overnight at RT in the dark. Whole-mount images were acquired using the Stellaris 8 Inverted Confocal Microscope (Leica) as previously described.

**RNA processing and Bulk RNA sequencing**

RNA was processed concurrently from eight patient organoid lines per diagnosis using the RNeasy Micro Kit (Qiagen, #74004). Briefly, matured spheroids from each line (n=8 per diagnosis) were filtered through a 100 µm strainer (Pluriselect), replated (15 spheroids/µL, 2:1 mixture of Matrigel and OWM) in two wells of a 12-well plate (3 domes/well) and allowed to stabilize overnight in 800 µL OGM. The following day, one well of spheroids/line was processed for RNA extraction, while the other well was switched to ODM for differentiation into colonoids for 3 days. For RNA extraction, the medium was removed, and Matrigel domes were dislodged with PBS from the bottom of the plate as previously described. To digest and remove any traces of Matrigel, the suspension was centrifuged (300 x g, 5 mins, 4^o^C), 350 µL of TrypLE Express (Gibco) was added to the spheroid pellets and incubated in a 37^o^C water bath for 3 mins. The tube was swirled briefly followed by the addition of 5 mL ice-cold OWM. Samples were washed (300 x g, 5 mins, 4^o^C), and the supernatant was aspirated. Any remnants of OWM were carefully removed with a 200 µL pipette and the resulting pellets were resuspended in the lysis buffer from the RNeasy Micro Kit and stored immediately at -80^o^C. Further processing of RNA followed the manufacturer’s instructions. RNA sequencing of poly-A enriched RNA libraries was carried out on the Illumina NovaSeq PE150 platform (Novogene). RNA-seq reads were aligned to the human reference hg38 genome and the nf-core/rnaseq (v3.5) workflow was used for developing the expression matrix.

**Ingenuity Pathway Analysis (IPA)**

Fold changes from the colonoid bulk RNA-seq data (Log2FC and p-adj values) were uploaded into the IPA software and used to predict canonical pathways that underlie transcriptional data. All upregulated and downregulated genes were concurrently analyzed in IPA with an expression p-value cutoff of 0.1. The top biological and cellular pathways and causal networks were selected based on the fisher’s exact test p-value which measures the overlap of observed and predicted regulatory gene sets.^5^

**Single Cell Data Analyses**

We utilized data from the established single-cell transcriptomic atlas of 68 colon biopsies obtained from 18 UC patients and 12 healthy controls ^6^. Processed single-cell expression matrices and metadata for human colon epithelial cells were obtained from the Single Cell Portal (<https://singlecell.broadinstitute.org/single_cell>, accession SCP259). Expression counts were normalized using the R (v.4.3.3) package Seurat (v.5.1.0). The reported PPARA, RXR, and lipid-related genes were visualized using the R package ComplexHeatMap (v.2.18.0).

**Quantitative RT PCR**

For most analyses, total RNA was processed from spheroids or colonoids by matrigel removal and digestion as previously described. Due to the short half-life of *UCP2*,^7^ samples for *UCP2* analysis were plated in a 1:1 mixture of Matrigel and OWM, differentiated, and lysed directly in the culture well without Matrigel removal. For all analyses, RNA was processed (RNeasy Micro Kit, Qiagen, #74004) and cDNA was synthesized using the High-Capacity cDNA Reverse Transcription kit (Applied Biosystems, #4368814). Gene expression was quantified on the QuantStudio 3 thermocycler using Taqman probes (table S2). Fold changes were quantified using the 2^- ΔΔCT^ method using *ACTB* or *GAPDH* as the reference gene where appropriate.

**LPL Activity**

Spheroids were plated, differentiated for 2 days or 5 days, and processed for Matrigel removal as previously described for RNA processing. Total colonoid protein was isolated using the Cell Lytic buffer (Sigma, #2978) with 1X Protease and Phosphatase inhibitor cocktail (ThermoFisher Scientific, #78440). Colonoids were incubated on ice for 30 mins with brief vortexing every 10 mins. Samples were centrifuged (10000 x g, 10 mins, 4^o^C), and the supernatant was transferred into fresh 1.5 mL tubes. Protein concentration was determined using the BCA protein Assay kit (ThermoFisher Scientific, #23227). LPL activity in each sample was determined in duplicates using the LPL activity assay kit (Cell Biolabs, #STA-610) following the manufacturer’s instructions, and the data was normalized to total protein concentration.

**Nuclear PPAR-α activity assay**

Three spheroids per group were differentiated in ODM for 24 hrs, pooled, and processed for Matrigel removal as previously described for RNA processing. Nuclei isolation followed the established Omni-ATAC protocol used for nuclei preparations for ATAC-seq.^8^ Nuclei protein concentration was determined using the BCA protein Assay kit (ThermoFisher Scientific, #23227). PPAR-α activity was assessed in the nuclear extracts with the PPAR-α activity kit (Abcam, #ab133107) following the manufacturer’s instructions.

**Extracellular Chemokine assay**

Spheroids were plated and differentiated in ODM for the indicated days in the figure legend, as previously described for RNA processing. ODM was collected at the indicated days and frozen at -80^o^C until analyses. For analyses involving differentiation until day 5, ODM was changed on day 2 before collection and analyses on day 5. The total colonoid protein was determined using the Cell Lytic buffer with 1X Protease and Phosphatase inhibitor as previously described. Extracellular chemokines in ODM on the indicated days were assessed using the following kits (CXCL1: Abcam #ab190805, CXCL11: ThermoFisher #EHCXCL11, CCL2: ThermoFisher #88-7399-22, CCL28: ThermoFisher #EHENC1) following manufacturer’s instructions.

**Statistical analyses**

Eight patient-derived colonoids were generated per diagnosis (control, active UC, inactive UC). For all data, symbols in figures represent the average measures of individual colonoid lines used for each assay. Analysis of two groups was conducted with a two-sided Mann-Whitney test or unpaired T-test. Wherever colonoid lines from the same diagnosis were treated with a Fccp, Fenofibrate, or GW6471, statistical analyses were done with a two-sided paired T-test or Wilcoxon signed-rank test. Two-way ANOVA with Tukey’s *post hoc* test was used for etomoxir experiments and those that were collected over 5 days (day 2 and day 5). Statistical analysis was conducted in Prism (v.10.0), and *P* values < 0.05 were considered statistically significant. For differentially expressed genes from the RNA-seq data, transcripts with Log_2_FC ≥ ±1, FDR P ≤ 0.1 were considered statistically significant.

| **Supplementary Tables and Figures**  **Supplementary Table 1. Participant demographic and clinical information** | | | | | | |
| --- | --- | --- | --- | --- | --- | --- |
|  | *n* | Control | *n* | Inactive UC | *n* | Active UC |
|  | 8 |  | 8 |  | 8 |  |
| Age, mean ± SD | 8 | 15.4 ± 2.5 | 8 | 14.1 ± 2.4 | 8 | 13.1 ± 3.2 |
| Sex, *n* (%) | 8 |  | 8 |  | 8 |  |
| Male |  | 2 (25) |  | 7 (88) |  | 4 (50) |
| Female |  | 6 (75) |  | 1 (13) |  | 4 (50) |
| Race, *n* (%) | 8 |  | 8 |  | 8 |  |
| African American |  | 2 (25) |  | 0 |  | 0 |
| Asian |  | 0 |  | 0 |  | 1 (12.5) |
| Caucasian |  | 6 (75) |  | 8 (100) |  | 7 (87.5) |
| Endoscopic Mayo Score, n (%) | 0 |  | 7 |  | 8 |  |
| 0 (Normal) |  | — |  | 7 (100) |  | 0 (0) |
| 1 (Mild) |  | — |  | 0 (0) |  | 1 (12.5) |
| 2 (Moderate) |  | — |  | 0 (0) |  | 6 (75) |
| 3 (Severe) |  | — |  | 0 (0) |  | 1 (12.5) |
| PUCAI Score, n (%) | 0 |  | 8 |  | 8 |  |
| 0 – 5 (Quiescent) |  | — |  | 8 (100) |  | 2 (25) |
| 10–30 (Mild) |  | — |  | 0 (0) |  | 3 (37.5) |
| 35–60 (Moderate) |  | — |  | 0 (0) |  | 3 (37.5) |
| 65-85 (Severe) |  | — |  | 0 (0) |  | 0 (0) |
| Disease Location, n (%) | 0 |  | 8 |  | 8 |  |
| Left-sided colitis |  | — |  | 1 (12.5) |  | 1 (12.5) |
| Extensive/Pancolitis |  | — |  | 7 (87.5) |  | 7 (87.5) |
| Treatment, n (%) | 0 |  | 8 |  | 7 |  |
| Oral 5-ASA |  | 0 (0) |  | 7 (88) |  | 5 (63) |
| Oral steroids |  | 1 (13) |  | 0 |  | 3 (38) |
| Rectal steroids |  | 0 (0) |  | 0 |  | 2 (25) |
| Anti-TNF biologic |  | 0 (0) |  | 2 (25) |  | 0 (0) |
| Infliximab |  | 0 (0) |  | 0 |  | 1 (13) |
| Vedolizumab |  |  |  | 0 |  | 1 (13) |

| **Supplementary Table 2. Taqman probes used for qPCR** | | | |
| --- | --- | --- | --- |
| **Manufacturer** | **Gene name** | **Species** | **Probe ID** |
| Invitrogen | *ACTB* | Human | Hs01060665_g1 |
| Invitrogen | *ASCL2* | Human | Hs00270888_s1 |
| Invitrogen | *ATOH1* | Human | Hs00944192_s1 |
| Invitrogen | *CCL2* | Human | Hs00234140_m1 |
| Invitrogen | *CCL28* | Human | Hs00219797_m1 |
| Invitrogen | *CHGA* | Human | HS00900370_m1 |
| Invitrogen | *CXCL1* | Human | Hs00236937_m1 |
| Invitrogen | *CXCL11* | Human | Hs00171138_m1 |
| Invitrogen | *FABP6* | Human | Hs01031183_m1 |
| Invitrogen | *GAPDH* | Human | Hs02758991_g1 |
| Invitrogen | *LGR5* | Human | Hs00969422_m1 |
| Invitrogen | *LPL* | Human | Hs00173425_m1 |
| Invitrogen | *MUC2* | Human | Hs00159374_m1 |
| Invitrogen | *UCP2* | Human | Hs01075227_m1 |
| Invitrogen | *WFDC2* | Human | Hs00196109_m1 |

**Supplementary Figures**

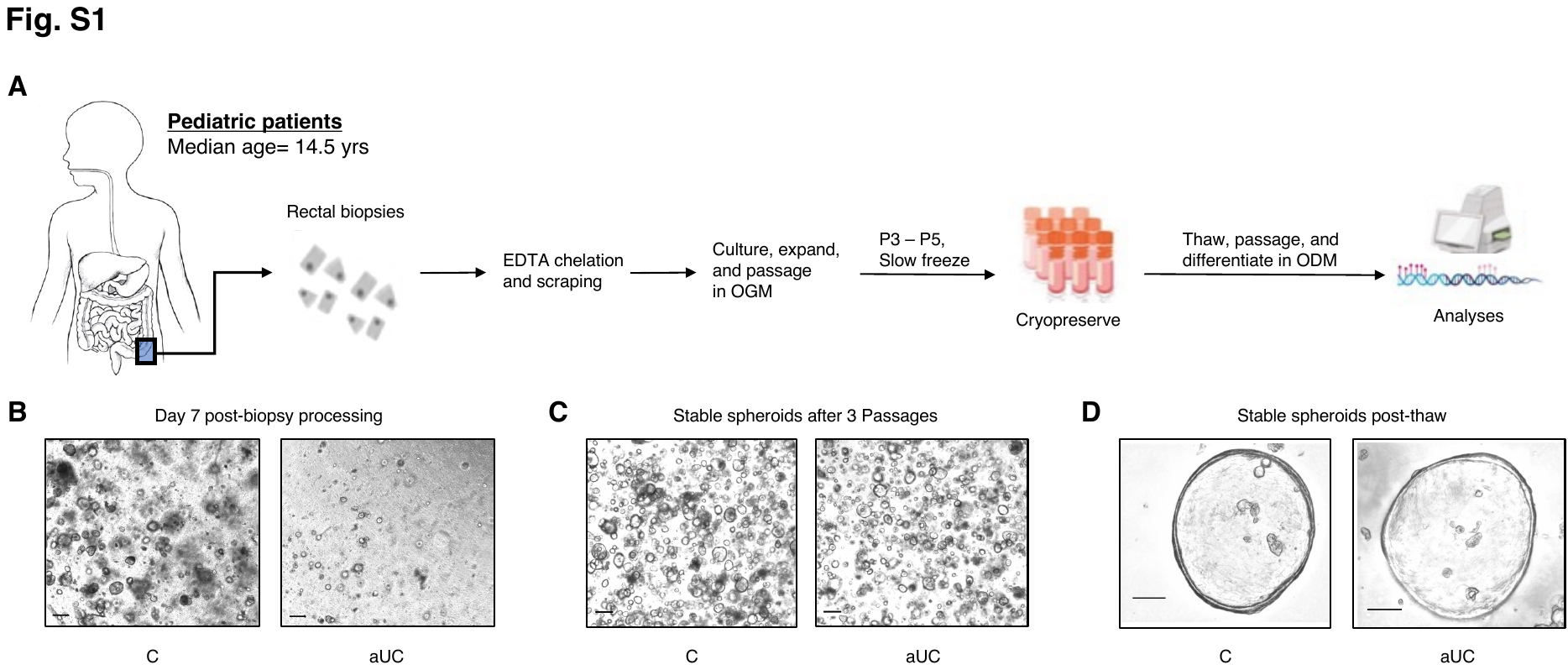

**Supplementary Figure 1. Colon Epithelial Spheroid Generation**

**(A**) Schematic of epithelial spheroids derivation and maintenance from pediatric rectal biopsies. Demographics of pediatric patients are provided in Supplementary Table 1. (**B**) Phase-contrast images of C and aUC spheroids after the first 7 days of biopsy derivation in OGM show that the development of aUC spheroids from biopsies is slower than the C spheroids. Scale bar, 200 µm. (**C**) Phase contrast images of C and aUC spheroids in culture after 3 passages, showing that after initial slow growth rates, aUC spheroids become more stable like C spheroids. Scale bar, 200 µm. (**D**) High magnification images of C and aUC spheroids, showing expected morphology with large lumens after freeze-thaw protocol. Scale bar, 50 µm. OGM, Organoid Growth Medium; ODM, Organoid Differentiation Medium

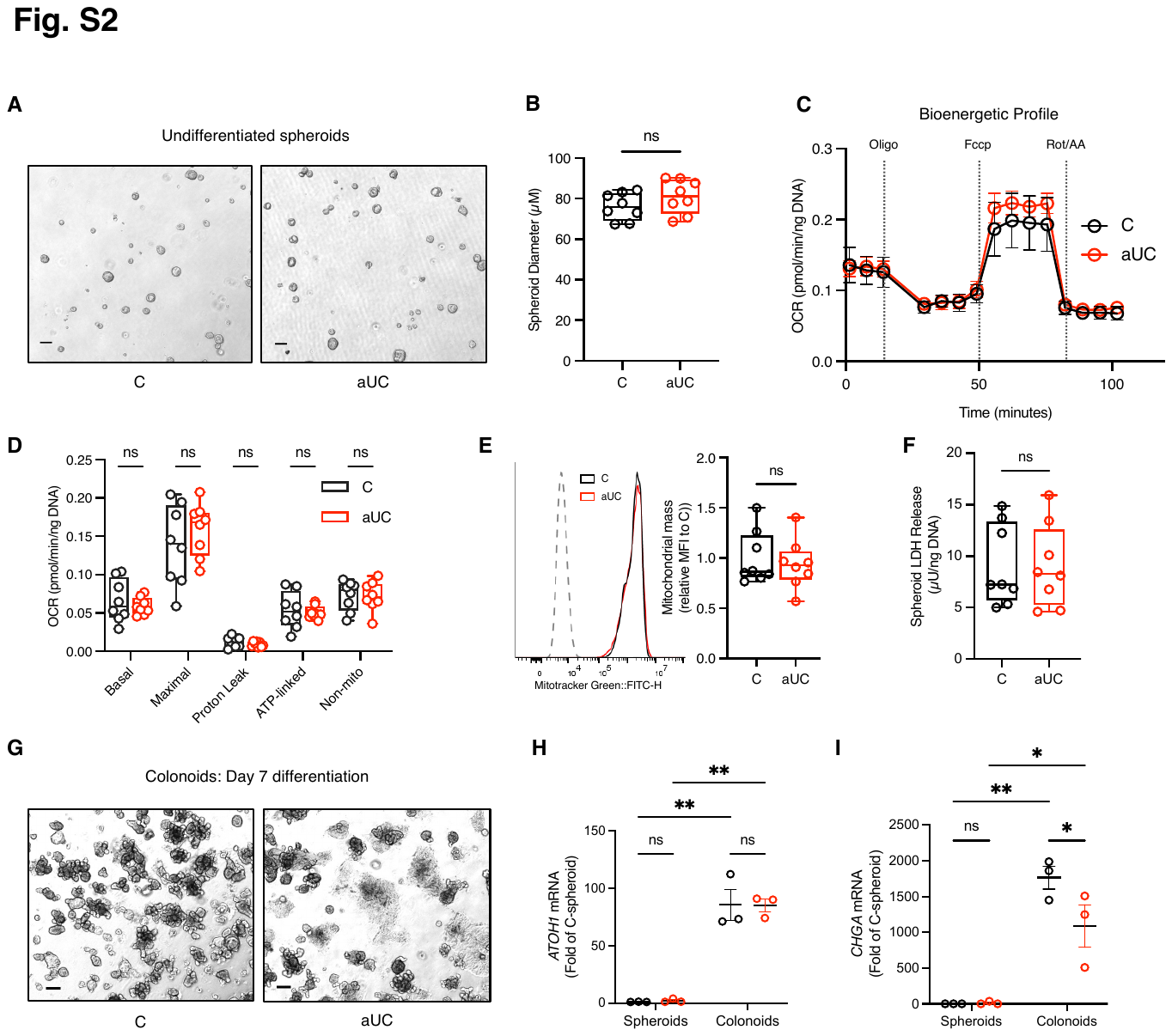

**Supplementary Figure 2. Pediatric colon spheroids exhibit similar morphological and metabolic profiles**

(**A**) Phase contrast images of undifferentiated C and aUC spheroids filtered after trypsinization, passed through a strainer and monitored for 7 days. Scale bar, 200 µm. (**B**) The diameter of C and aUC spheroids described in **A.** Blinded measures of the diameter of all spheroids on a high-powered field were taken. n=8 patient spheroid line/group, two-sided Mann-Whitney test. (**C**) Bioenergetic profile of C and aUC spheroids from Seahorse MitoStress test. n=8 patient spheroid line/group. (**D**) OCR of C and aUC spheroids cultured in unstimulated conditions in OGM. n=8 patient spheroid line/group, unpaired, two-sided t-test. (**E**) Estimation of mitochondrial mass with Mitotracker green intensity in undifferentiated C and aUC spheroids. n=8 patient spheroid line/group, two-sided Mann-Whitney test. (**F**) LDH activity in C and aUC spheroids in the medium from the same plate as (**C** and **D**). n=8 patient spheroid line/group, unpaired, two-sided t-test. (**G**) Phase contrast images of C and aUC differentiated spheroids (colonoids) on day 7 of differentiation. Scale bar, 200 µm. (**H**) Spheroids embedded in Matrigel were cultured in OGM and paired samples were differentiated in ODM for 3 days (colonoids) and assay for (**H**) *ATOH1* and (**I**) *CHGA* mRNA abundance using qRT-PCR. n=3 patient line/group, 2-way ANOVA. *=*P* < 0.05, **=*P* < 0.01. For all data, symbols in boxplots represent the average measures of individual spheroid lines. OCR, oxygen consumption rates; ODM, organoid differentiation medium; OGM, organoid growth medium

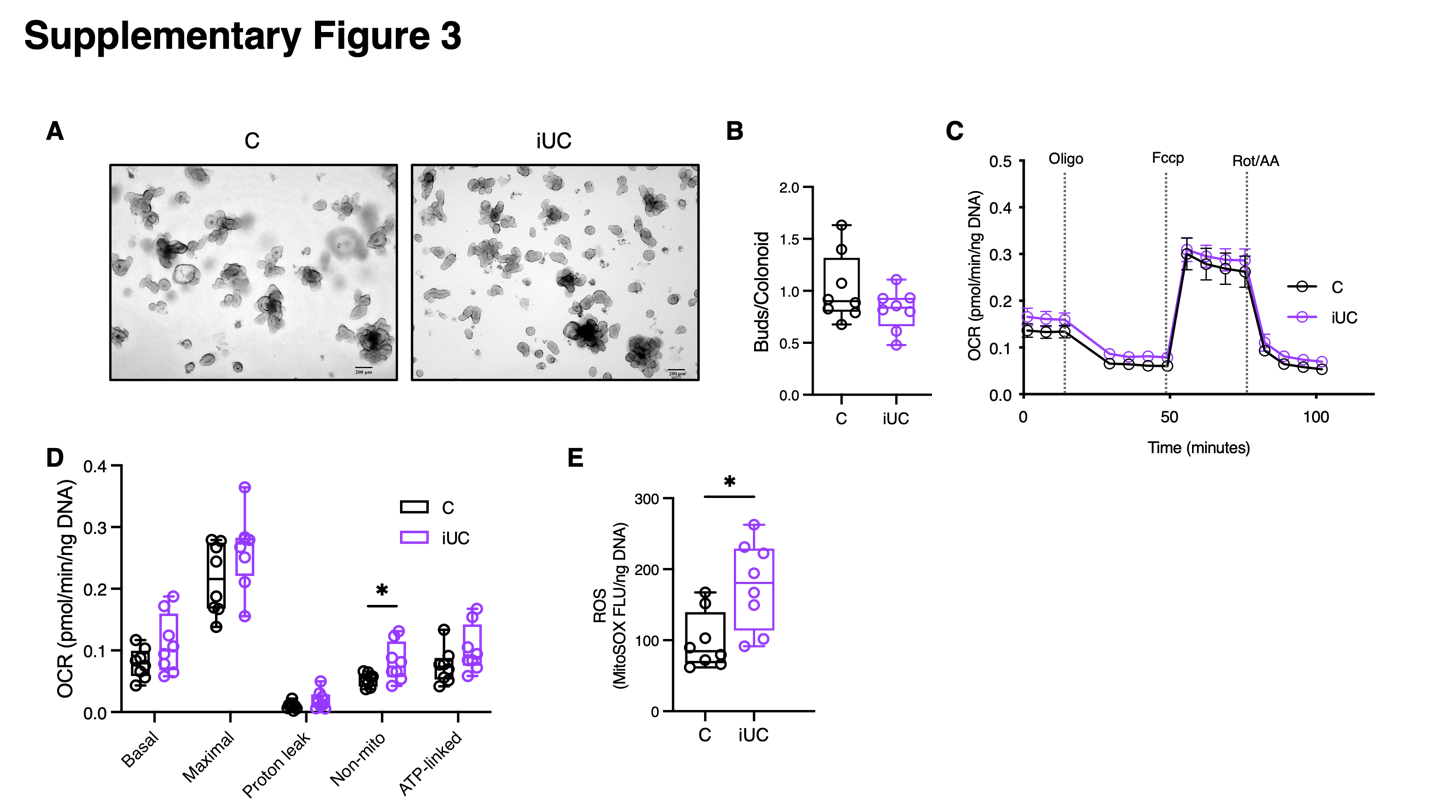

**Supplementary Figure 3. Morphologic and Metabolic similarities in inactive UC and control colonoids**

(**A**) Phase contrast images of pediatric C and iUC colonoids after 3-day differentiation in unstimulated conditions in ODM. Scale bar, 200 µm. (**B**) The number of buds/colonoid in colonoids treated as in **a**. n=8 patient colonoid line/group, two-sided Mann-Whitney test. (**C**) Bioenergetic profile of C and iUC colonoids differentiated for 3 days as in **A**, and subjected to the Seahorse MitoStress test. n=8 patient colonoid line/group. (**D**) MitoStress OCR response of C and iUC colonoids differentiated for 3 days as in **A**, in 96-well Seahorse plate. n=8 patient colonoid line/group, Mann-Whitney test. (**E**) The same cohort of colonoids cultured and used in **D**, were used to estimate mitochondrial ROS with the MitoSOX fluorescence assay. n=8 patient colonoid line/group, two-sided Mann-Whitney test. For all data, symbols in boxplots represent the average measures of individual colonoid lines. *=*P* < 0.05, **=*P* < 0.01, ***=*P* < 0.001. OCR, oxygen consumption rates; ODM, organoid differentiation medium; OGM, organoid growth medium; ROS, reactive oxygen species

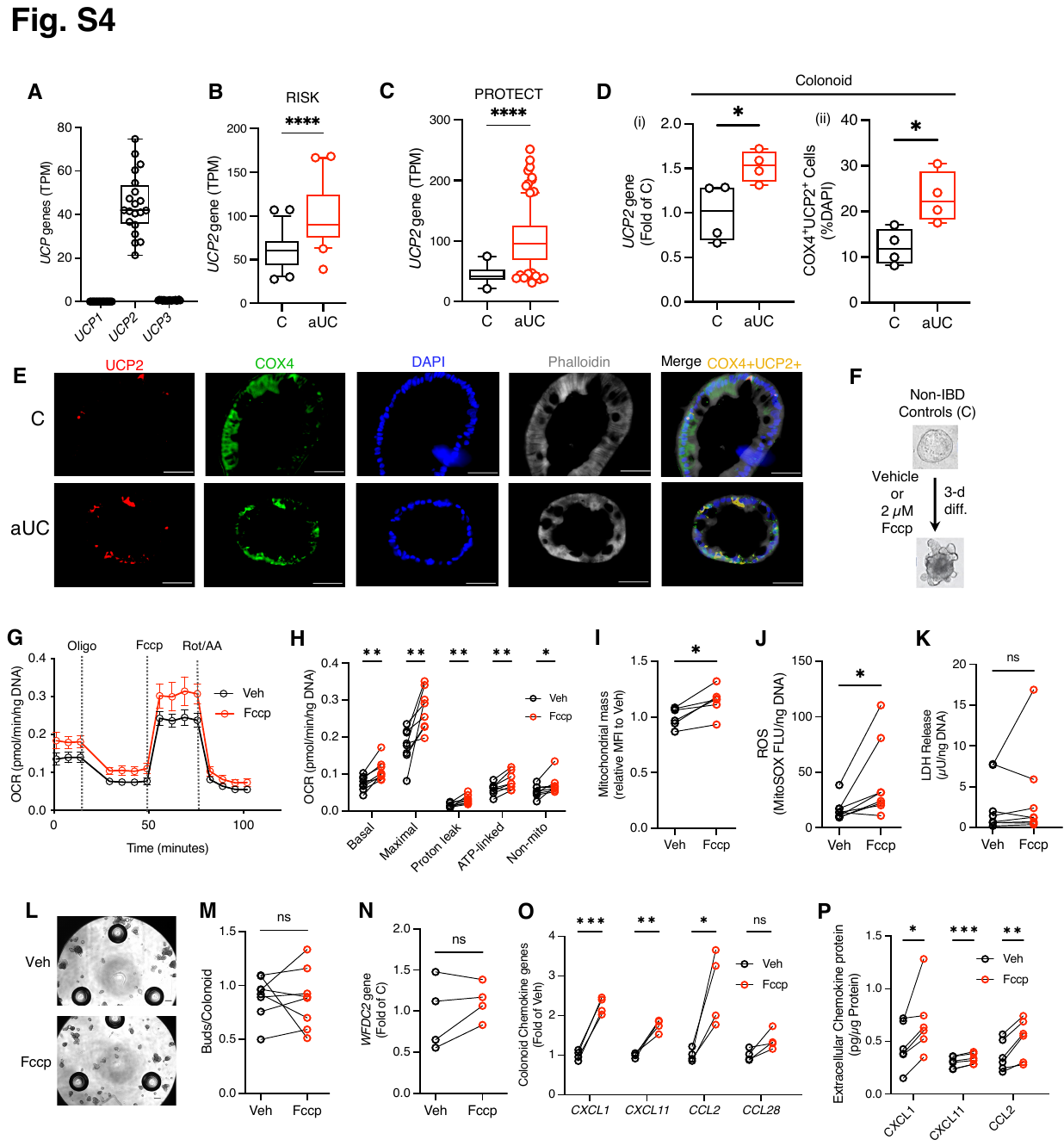

**Supplementary Figure 4. Metabolic uncoupling contributes to hypermetabolic phenotypes during colonoid differentiation**

(**A**) Rectal gene expression of uncoupling proteins in bulk RNA-Seq data from non-IBD control participants in the NIH PROTECT study showing dominant UCP2 gene expression over other uncouplers (transcripts per million (TPM) values). n=20. (**B**) Rectal UCP2 gene expression (TPM) from treatment-naïve participants with UC in the RISK cohort. n=43-55/group. Two-tailed Mann-Whitney test. (**C**) Rectal UCP2 gene expression (TPM) from the NIH PROTECT study of treatment-naive pediatric UC patients. n=20(C) -206(aUC) /group. Two-tailed Mann-Whitney test. (**D**) (i) UCP2 gene expression assessed by qRT-PCR and (ii) UCP2 protein expression in pediatric C and aUC colonoids differentiated for 3 days in ODM. n=4 patient colonoid line/group i) two-tailed unpaired t-test. ii) two-sided Mann-Whitney test. (**E**) C and aUC colonoids were embedded in OCT after 3-day differentiation. Representative immunofluorescence images of C and aUC colonoids after 3-day differentiation showing UCP2 expression with the mitochondrial marker, COX4. Semi-quantitation of COX4+UCP2+ cell counts is presented in **D**(ii). Scale bar, 50 µm. (**F**) Morphological and metabolic effects of mitochondrial uncoupling during colonoid differentiation were tested in C colonoids treated with or without the mitochondrial protonophore, FCCP. (**G**) Bioenergetic profile of C colonoids treated as in **F**, and subjected to the Seahorse MitoStress test. n=8 patient colonoid lines/group. (**H**) MitoStress OCR response of C colonoids treated as in **F**, in 96-well Seahorse plate. n=8 patient colonoid line/group, two-sided paired t-test. (**I**) Estimation of mitochondrial mass with Mitotracker green intensity in C colonoids treated as in **F**. n=8 patient spheroid line/group, two-sided paired t-test. (**J**) Mitochondrial ROS with the MitoSOX fluorescence assay in C colonoids treated as in **F**. n=8 patient colonoid line/group, two-sided Wilcoxon signed-rank test. (**K**) LDH activity in the medium after differentiation for 2 days. n=8 patient colonoid lines/group, two-sided Wilcoxon signed-rank test. (**L**) Representative phase contrast images of pediatric C colonoids treated as in **F**. Scale bar, 200 µm. (**M**) The number of buds/colonoids in colonoids treated as in **F**. n=8 patient colonoid line/group, two-sided Wilcoxon signed-rank test. (**N**) WFDC2 gene expression in colonoids treated as in **F**, assessed by qRT-PCR. n=4 patient colonoid lines/group, paired t-test. (**O**) Gene expression of chemokines in colonoids treated as in **F**, assessed by qRT-PCR. n=4 patient colonoid line/group, paired t-test. (**P**) Colonoids were treated as in **F**, and Chemokine protein concentrations were assessed in ODM after a 2-day differentiation. n=6 colonoid lines/group, paired t-test. For all data, symbols represent the average measures of individual spheroid lines. *=*P* < 0.05, **=*P* < 0.01, ***=*P* < 0.001. LDH, lactate dehydrogenase; OCR, oxygen consumption rates; ODM, organoid differentiation medium; ROS, reactive oxygen species

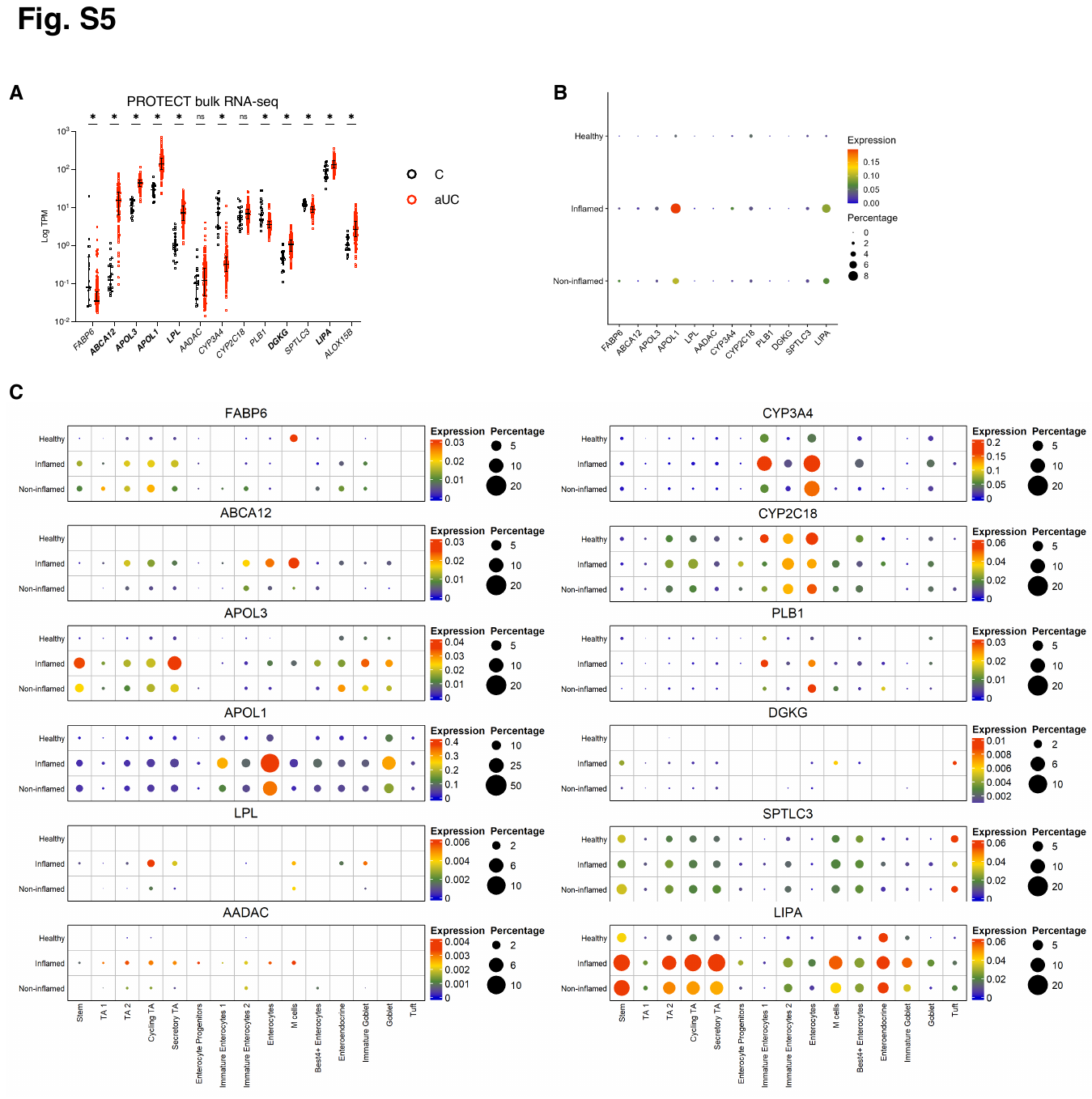

**Supplementary Figure 5. Dysregulated lipid metabolism genes in the colon epithelia of active UC**

(**A**) Rectal gene expression (TPM) of lipid-related colonoid genes (as in Fig. 4**A**) from the NIH PROTECT study ^9^ of treatment-naive pediatric UC patients. n=20(C) -206(aUC)/group. Kruskal-Wallis test with Dunn’s multiple comparisons. (**B**) Expression of colonoid lipid-related genes (as in Fig. 4A) by disease state in publicly available adult single-cell (sc) RNA-seq data ^6^. Data is colored by relative gene expression and the relative size of each dot shows the percentage of cells expressing each marker per disease state. (**C**) Expression of colonoid lipid-related genes (as in Fig. 4A) in scRNA-seq data ^6^ by epithelial cluster. Data is colored by relative gene expression and the relative size of each dot shows the percentage of cells expressing each marker per epithelial cluster.*=*P* < 0.05.

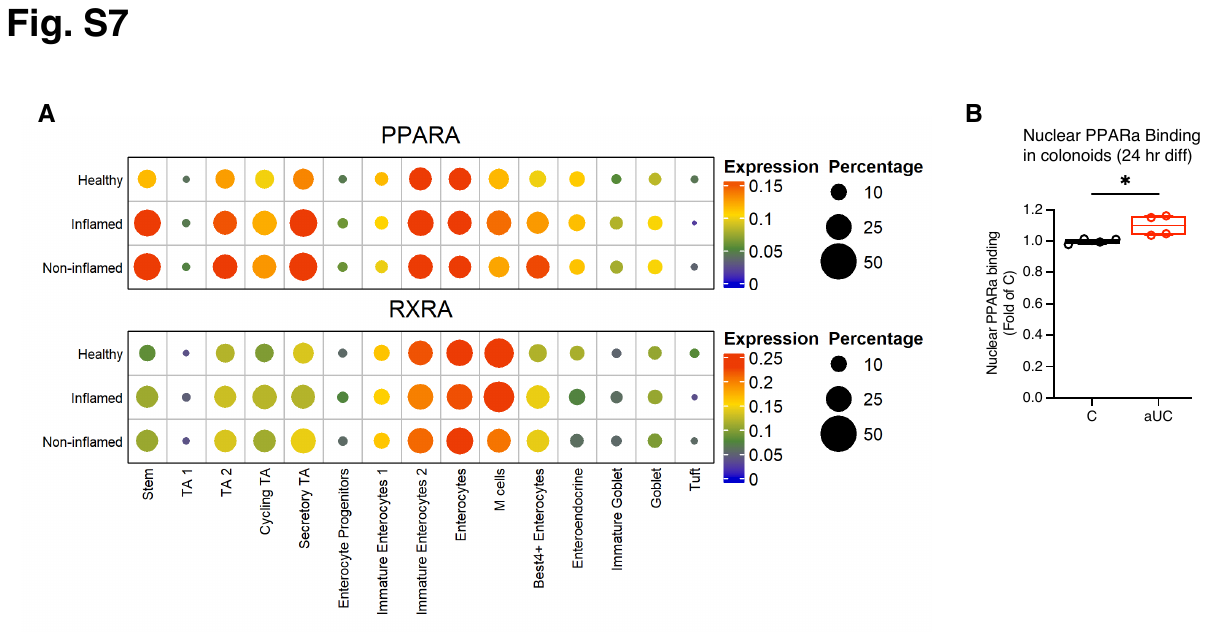

**Supplementary Figure 6. Epithelial PPARA expression in active UC**

(**A**) Colon scRNA-seq data ^6^ showing the expression of PPARA and RXRA by disease state in colon epithelial lineages. Data is colored by relative gene expression and the relative size of each dot shows the percentage of cells expressing each marker per disease state. (**B**) PPAR-α activity assay in nuclear extracts of pooled C and aUC colonoids (n=3/group) differentiated for 24 hrs. Symbols represent four technical replicates from pooled nuclear extracts and are presented relative to C. Two-sided unpaired t-test. *=*P* < 0.05.

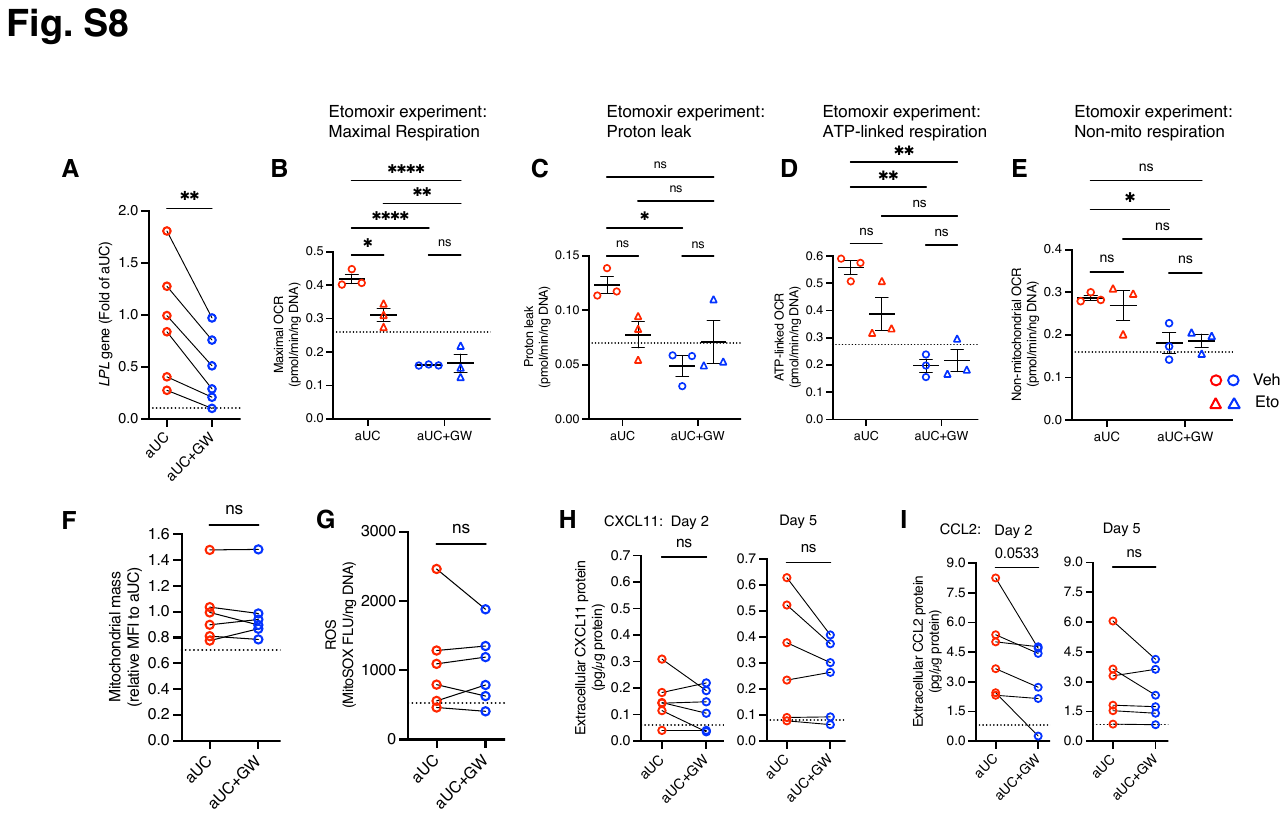

**Supplementary Figure 7. Blockade of lipid synthesis suppresses hypermetabolic features in active UC colonoids**

aUC colonoids were treated with 1 µM of the PPAR-α antagonist, GW6471 (GW), or vehicle (EtOH) during differentiation. (**A**) Effect of GW on *LPL* gene expression by qRT-PCR. n=6 patient colonoid lines/group, two-sided paired t-test. (**B**) Maximal, (**C**) Proton leak, (**D**) ATP-linked, and (**E**) Non-mitochondrial OCR response of aUC colonoids differentiated for 3 days in ODM with or without GW and subjected to the Seahorse MitoStress test in a nutrient-deprived medium with or without Etomoxir. n=3 patient colonoid lines/group. 2-way ANOVA, Tukey’s *post hoc* test. (**F**) Estimation of mitochondrial mass with Mitotracker green intensity in aUC colonoids treated with or without GW in ODM for 3 days. n=6 patient spheroid line/group, two-sided Wilcoxon signed-rank test. (**G**) ROS estimate with the MitoSOX fluorescence assay in a UC colonoids differentiated for 3 days with or without GW. n=6 patient colonoid line/group, two-sided Wilcoxon signed-rank test. (**H)** aUC colonoids with or without GW were differentiated in ODM for 5 days. ODM collected on day 2 and on day 5 (representing the last 3 days of differentiation) was used to assess CXCL11 and (**I**) CCL2 secretion. n=6 patient colonoid lines/group, two-sided paired t-test. For all data, symbols represent the average measures of individual colonoid lines. *=*P* < 0.05, **=*P* < 0.01, ***=*P* < 0.001. ****=*P* < 0.0001 OCR, oxygen consumption rates; ODM, organoid differentiation medium; MFI, mean fluorescence intensity
